# Single-cell splice isoform usage reveals distinct axes of cellular identity and senescence

**DOI:** 10.64898/2026.09.11.748700

**Authors:** Madhav Mantri, Angela M. Detweiler, Jaeyoon Lee, Jamie Ha-Young Kwon, Andy Zhou, Laura Tong, Robert C. Jones, Norma F. Neff, Tabula Sapiens Consortium, Stephen R. Quake

## Abstract

Alternative splicing greatly expands the diversity of gene products encoded by the human genome. Single-cell transcriptomic atlases have characterized human cell types through gene-level expression, but short-read sequencing has limited the ability to resolve full-length isoforms and their functional consequences. Here, we present a cross-tissue single-cell long-read isoform atlas spanning 26 human tissues. We identify hundreds of thousands of novel isoforms along with their cell-type-specific usage, and discover that over one-third of expressed isoforms are absent from existing reference databases. We further demonstrate that isoform usage is a structured, measurable axis of cellular identity that is distinct from gene expression. Applying this framework to cellular senescence, we resolve p16^INK4a^ and p14^ARF^ transcripts from the *CDKN2A* locus in individual cells and uncover cell-type-dependent isoform remodeling associated with the p16^INK4a^ senescence program. This isoform-resolved single-cell atlas offers a versatile framework to dissect the cellular logic of isoform regulation in senescence and beyond.

## Introduction

Alternative splicing is a major source of transcript and protein diversity, with alternative splice products generated from most human genes^1–4^. Single-cell transcriptomic atlases, including Tabula Sapiens and the Human Cell Atlas, have defined human cell types and states at organism scale from gene-level expression^5–12^. However, conventional 3’- and 5’-end short-read sequencing captures only fragments of individual transcripts, making full-length isoform assignment impossible. Although junction-based approaches such as SICILIAN and SpliZ enable cell-resolved inference of splicing from short reads^13,14^, they cannot reconstruct complete transcript structures, coding frames, or untranslated regions. Thus, the extent to which isoform usage contributes independently to cell identity and state remains largely unresolved.

Full-length single-cell long-read sequencing directly addresses this limitation by reading individual barcoded cDNA molecules end to end, thereby enabling direct measurement of complete transcript structures, including alternative exons, coding sequences, and UTRs. Recent high-throughput approaches, such as MAS-ISO-Seq^15^, have substantially increased the number of full-length molecules recovered per run, making large-scale single-cell long-read atlases feasible^16–23^. Such datasets provide an opportunity to ask whether isoform usage encodes cell identity beyond gene-level expression, whether isoform programs are organized primarily by cell type or tissue of residence, to what extent previously unannotated isoforms are recurrently expressed across tissues and cell types, and what consequences isoform variation has for coding potential, nonsense-mediated decay (NMD)^24^, and transcript boundaries.

These questions are particularly relevant to cellular senescence, in which alternative splicing is progressively remodeled during aging and can actively reinforce cell-state transitions^25–27^. A prominent example is the *CDKN2A* locus, which produces the distinct tumor suppressors p16^INK4a^ and p14^ARF^ through alternative first exons and reading frames^28,29^. p16^INK4a^ acts through Rb to enforce the stable proliferative arrest characteristic of senescence^30^, whereas p14^ARF^ activates p53 across a broader range of stress responses^31–33^. Whether individual senescent cells express p16^INK4a^, p14^ARF^, or both cannot be resolved from gene-level measurements and requires direct identification of transcript isoforms.

Here, we present a cross-tissue single-cell long-read isoform atlas of human tissues comprising 60 libraries from 26 tissues and 12 donors. After unifying transcript identifiers across samples, we obtained 203,311 cells spanning 144 annotated cell types and 854,410 robustly detected transcript structures. Of these, 37.6% are novel structures; these structures recur across independent samples and donors and are supported by matched short-read data at both splice-junction and single-cell levels. We use this atlas to show that isoform usage captures cell identity beyond gene-level expression, to partition isoform variation between cell type and tissue of residence, and to define consequences for coding potential, NMD, and transcript boundaries. Finally, we resolve p16^INK4a^ and p14^ARF^ transcripts in individual cells and identify isoform-level remodeling associated with the p16^INK4a^ senescence program.

## Results

### A cross-tissue single-cell atlas of full-length human transcript isoforms

We generated full-length single-cell transcriptomes from 26 tissues collected across 12 Tabula Sapiens donors using long-read RNA-seq chemistry (**Methods, Fig. 1A–B**). We sampled at least two donors per tissue, including one male and one female donor for each non-reproductive tissue except ear and pancreas, which were sampled from male donors only. We processed raw long reads through a unified pipeline to generate isoform-resolved single-cell gene expression profiles for 60 3’ scRNA-seq samples (**Methods; Supp. Fig. 1A**). Isoform identifiers were harmonized across tissues and donors by a global re-collapse, enabling direct atlas-wide comparisons of isoform usage (**Methods**; **Supp. Fig. 1B**). After quality control filtering, we analysed long-read single-cell transcriptomes for 203,311 cells across 26 tissues and organs (**Supp. Fig. 1C**).

**Figure 1.**
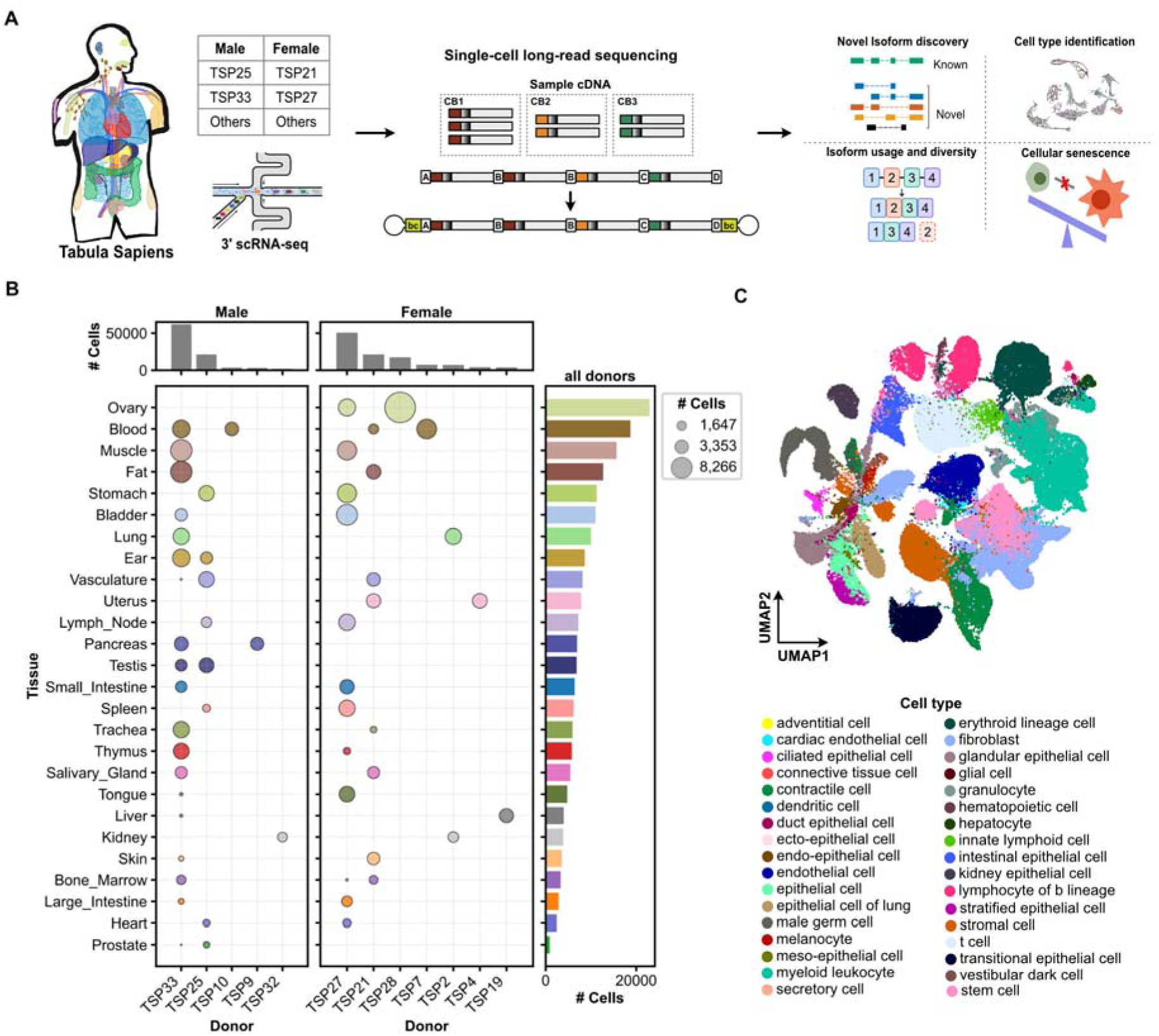
A cross-tissue single-cell atlas of full-length human transcript isoforms. **(A)** Experimental overview. Tissues were dissociated from Tabula Sapiens donors and processed by 10x Genomics 3’ scRNA-seq, generating cDNA in which each molecule carries a cell barcode and a UMI. cDNA was concatenated into multi-segment molecules with segment adapters separating individual cDNAs and a sample barcode (bc) at each end to generate long-read RNA-seq data. Concatenated reads were subsequently deconcatenated and demultiplexed to recover single-cell, full-length isoform data. **(B)** Cells recovered per donor and tissue. Dot plots show the number of cells per donor–tissue combination for male (left, 5 donors) and female (middle, 7 donors) donors; dot area is proportional to cell number and color encodes tissue. Top marginal bars give the total cells per donor. Right, total cells per tissue summed across all 12 donors. Donors are Tabula Sapiens (TSP) identifiers, and tissues are ordered by total cell number. **(C)** UMAP embedding of all profiled single cells, clustered by isoform expression and colored by broad cell class annotation (legend at bottom).

The cells in each of the 26 tissues and organs were annotated by consensus prediction (popV) and manually renamed in accordance with the Cell Ontology, resulting in 144 precisely defined cell types grouped into 34 broad cell classes. Dimensionality reduction on the most variable isoform features separated cells by broad cell class (**Fig. 1C**). Since we split the cDNA from the same dissociated cell suspensions and processed them with both 3’ short-read and long-read RNA sequencing, the resulting long-read datasets could be matched to existing short-read scRNA-seq datasets from the same individual cells in the Tabula Sapiens atlas. Of the 197,994 cells from the 58 libraries with a matched short-read library, 160,891 (81.3%) were also detected in the short-read data, corresponding to 79.1% of the full 203,311-cell atlas. Thus, most long-read cells have a short-read counterpart available for direct comparison (**Supp. Fig. 2A**). We then restricted the analysis to these shared cells and generated donor-level pseudobulk gene expression profiles. Gene expression measurements showed strong agreement between platforms, with a mean Pearson correlation of 0.81 across donors (range 0.71–0.89) and a mean Spearman correlation of 0.82 (range 0.79–0.84) (**Supp. Fig. 2B–C**). For reference, two replicate long-read libraries prepared from the same tissue agreed at r = 0.945 and ρ = 0.894 (**Supp. Fig. 2D**). These results demonstrate that full-length single-cell long-read sequencing accurately quantifies gene expression while providing isoform-level information that short-read sequencing alone cannot resolve.

**Figure 2.**
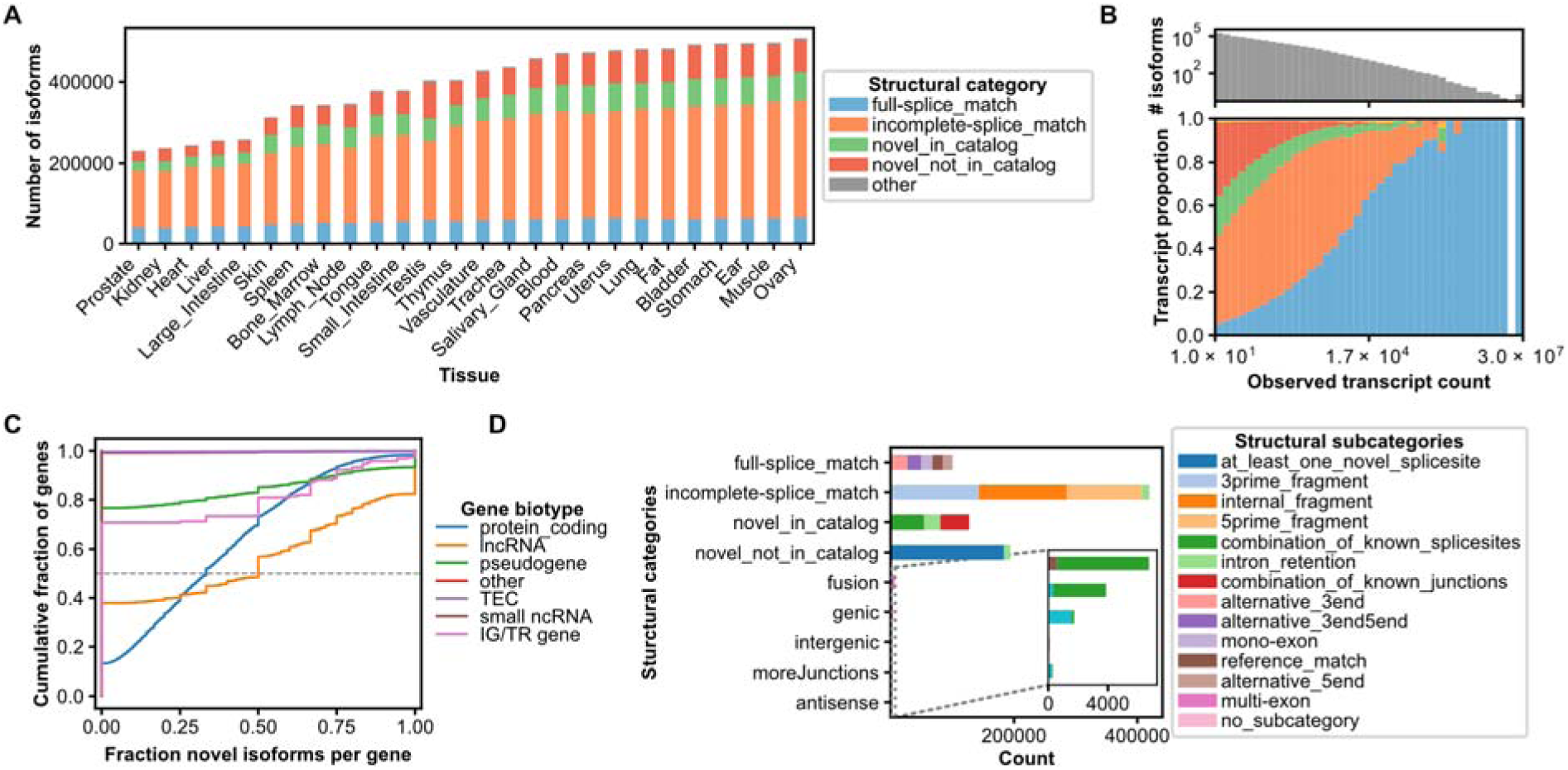
Novel isoforms reveal extensive transcript structural diversity across human tissues. **(A)** Number of isoforms recovered per tissue, colored by structural category relative to the GENCODE reference: full-splice match (FSM), incomplete-splice match (ISM), novel-in-catalog (NIC), novel-not-in-catalog (NNC), and others (fusion, genic, intergenic, moreJunctions, and antisense). Tissues are ordered by total isoform count. **(B)** Isoform abundance and structural composition as a function of expression. Top: number of isoforms (log scale) per observed transcript-count bin. Bottom: proportion of isoforms in each structural category across the same bins (colors as in A). **(C)** Empirical cumulative distribution of the fraction of a gene’s expressed isoforms that are novel, stratified by gene biotype. The dashed grey line marks the 0.5 quantile. **(D)** Counts of isoforms per structural category, decomposed by structural subcategory — including at least one novel splice site, combinations of known splice sites/junctions, internal/5’/3’ fragments, intron retention, mono- and multi-exon forms, alternative 5’/3’ ends, and reference match.

### Widespread novel isoforms expand the human transcript landscape

We first characterized the structural composition of isoforms across the atlas at genome-wide scale. Isoforms were classified as transcripts that exactly match a previously annotated splice junction chain (full-splice-match, FSM), those that match only a subset of a previously annotated junction chain due to 5’ or 3’ variation or intron retention (incomplete-splice-match, ISM), those that contain a novel combination of known splice sites (novel-in-catalog, NIC), those that contain at least one previously unannotated splice site (novel-not-in-catalog, NNC), and “Other” (transcripts falling outside a single annotated gene model). (**Fig. 2A**). We applied this analysis to isoforms containing at least 10 unique molecules across at least five cells to distinguish robust transcript structures from low-confidence events. Of 854,410 distinct transcript structures, only 11.8% were FSMs, whereas 49.1% were ISMs. Notably, more than one-third of the robustly detected transcript structures were novel, comprising 22.7% NNC and 14.9% NIC isoforms (**Supp. Fig. 3A**), closely matching independent long-read surveys^22,23^. “Other” transcripts included those joining two or more genes (fusions), transcripts spanning multiple genes with shared junctions (moreJunctions), transcripts overlapping a gene on the same strand while retaining intronic sequence (genic), transcripts overlapping genes only on the opposite strand (antisense), and transcripts falling outside annotated genes (intergenic); these accounted for less than 2% of all structures.

**Figure 3.**
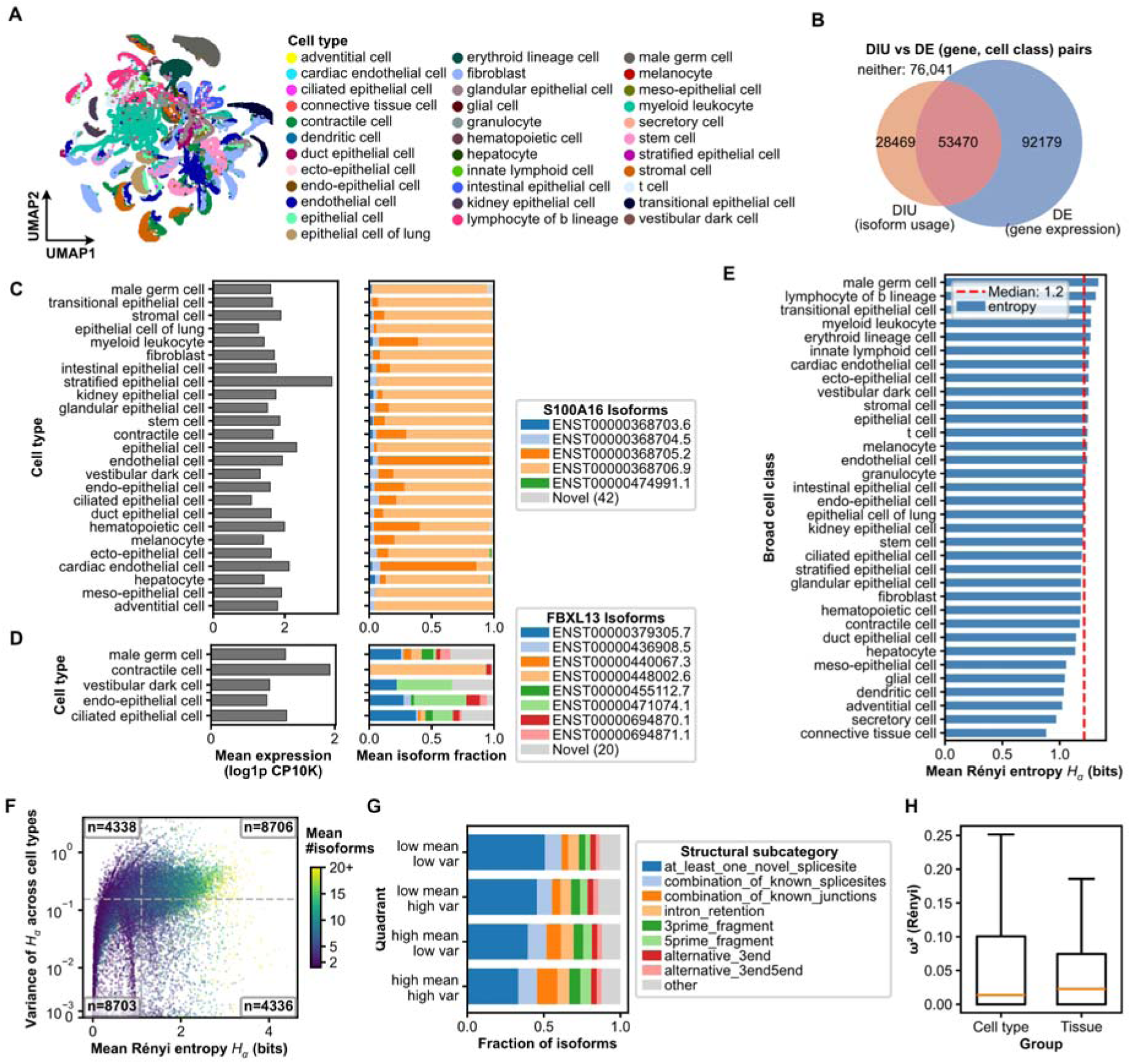
Isoform usage defines cellular identity beyond gene expression. **(A)** UMAP embedding of 203,311 cells computed from isoform fractions alone, colored by broad cell class annotation. **(B)** Overlap between the set of genes with differential isoform usage and the set of differentially expressed genes, across all gene and cell class pairs testable on both axes. Regions are area-proportional; pairs significant on neither axis are not drawn and are reported separately. **(C)** Gene expression and isoform usage of S100A16 across broad cell classes. Isoform usage is shown as the mean fraction of the gene’s molecules assigned to each isoform. Novel isoforms are grouped into a single category. A cell class is shown only if at least 5 of its cells express the gene and those cells make up at least 1% of the class; classes failing either threshold are omitted from both panels. **(D)** Gene expression and isoform usage of FBXL13 across broad cell classes. Isoform usage is shown as the mean fraction of the gene’s molecules assigned to each isoform. Novel isoforms are grouped into a single category. A cell class is shown only if at least 5 of its cells express the gene and those cells make up at least 1% of the class; classes failing either threshold are omitted from both panels. **(E)** Mean Rényi entropy Hα of isoform usage within each broad cell class, ordered by decreasing entropy. The dashed red line marks the median across classes. **(F)** Per-gene mean Rényi entropy across cell types plotted against the variance of that entropy across cell types (log scale), colored by the mean number of isoforms detected per gene. Dashed lines mark the median of each axis, and quadrant labels give the number of genes falling in each. Restricted to the 26,083 genes tested in at least two cell types. **(G)** Structural subcategory composition of the isoforms in each mean-versus-variance quadrant of (F), as a fraction of that quadrant’s isoforms. Quadrants are ordered by increasing mean entropy. **(H)** Distribution of df-adjusted ω^2^ for isoform diversity attributable to cell type and to tissue. Boxes span the interquartile range, orange lines mark medians, and whiskers extend to 1.5 times the IQR.

Across 60 samples, we recovered 520,216 known isoforms (FSM and ISM) and 321,443 novel isoforms (NIC and NNC). Novel isoforms were broadly reproducible across samples, with 97.1% detected in at least two samples and a median of 10 samples per isoform, comparable to the 98.8% of known structures detected across multiple samples (**Supp. Fig. 3B**). Novel isoforms were nevertheless less abundant and detected in fewer cells than known isoforms (**Supp. Fig. 3C**). Novel categories were enriched among lower-abundance transcripts, whereas FSMs dominated the highly expressed range (**Fig. 2B**). ISMs were shorter than FSMs but contained more exons (**Supp. Fig. 3D**), reflecting the enrichment of FSMs for mono-exon transcripts and the predominance of multi-exon structures among ISMs.

Isoform repertoires are widespread across genes. Expressed genes contained a median of six isoforms, with 80.6% expressing multiple isoforms, consistent with the bulk literature (**Supp. Fig. 3E**). Novel isoforms were detected across gene biotypes, with 64% of expressed genes producing at least one novel isoform, although their contribution varied across biotypes (**Fig. 2C; Supp. Fig. 3F**). Despite their broad distribution across genes, novel isoforms accounted for only 5.4% of detected molecules (**Supp. Fig. 3G**). Thus, although individual novel isoforms were generally lowly expressed, transcript diversity was widespread across the genome, with most expressed genes producing multiple isoforms and a substantial fraction producing previously unannotated structures.

Full-length sequencing further revealed substantial structural variation within annotated isoform classes. Among 82,428 multi-exon FSMs, only 20.6% matched a reference transcript at both ends; 33.2% used an alternative 3’ end, 19.0% an alternative 5’ end, and 27.1% differed at both ends (**Fig. 2D**). A further 101,111 NIC isoforms were built from new combinations of annotated splice sites or junctions (**Fig. 2D**). Retained introns were present in 49,587 isoforms spanning several structural categories, including the remaining 26,322 NIC structures (**Fig. 2D**). Thus, matching an annotated splice-junction chain did not necessarily imply recovery of the annotated isoform, as reported previously across human tissues^34^. These structural differences were associated with substantial differences in predicted transcript fate. Predicted NMD targets comprised 6.3% of coding isoforms and were enriched among novel isoforms, reaching 11.7% of all NIC and 7.7% of all NNC isoforms compared with 2.6% of FSMs and 0.9% of ISMs (**Supp. Fig. 3H**). Coding potential varied markedly across isoform classes, ranging from 87.3% in moreJunctions and 79.0% in ISMs to 17.3% in intergenic and 13.2% in antisense isoforms, with 69.6% of expressed isoforms predicted to encode proteins (**Supp. Fig. 3I**). Together, these results show that transcript structural diversity is associated with distinct predicted coding and post-transcriptional fates. These structural and functional features varied substantially across tissues and cell types, revealing cell-type-specific differences in isoform architecture and coding potential (**Supp. Fig. 4A–B**).

**Figure 4.**
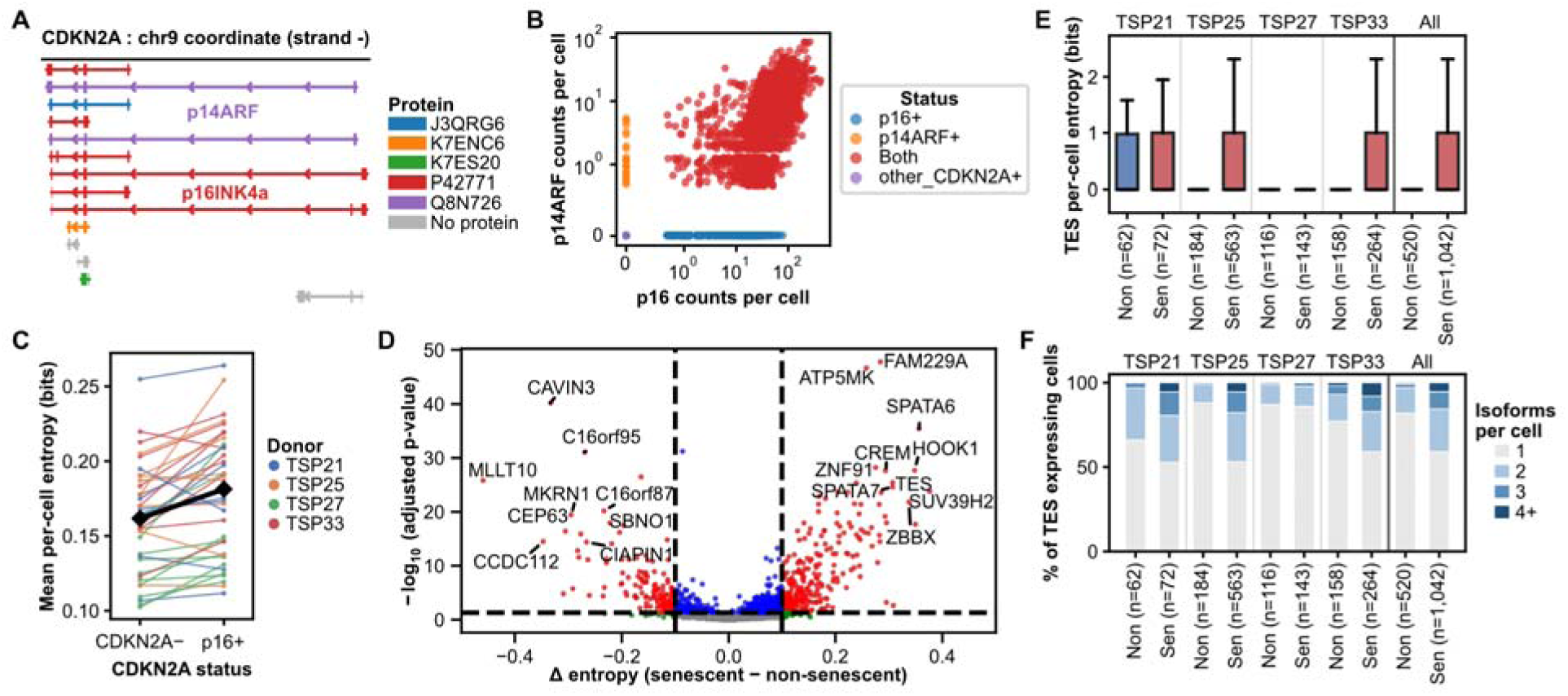
The p16^INK4a^ senescence program is associated with widespread remodeling of transcript isoform usage. **(A)** Structure of the *CDKN2A* locus. Alternative first exons and reading frames produce two distinct products: p16^INK4a^ (6 annotated transcripts) and p14^ARF^ (2 annotated transcripts), shown as full-length isoform models. **(B)** Per-cell p16^INK4a^ versus p14^ARF^ counts from the targeted *CDKN2A* capture, on a symlog scale with integer counts jittered for visibility. Cells are colored according to *CDKN2A* status (p16^INK4a^-positive, p14^ARF^-positive, both, other *CDKN2A*-positive). **(C)** Per-cell isoform entropy by sample, on library-size-matched cells. Entropy is averaged over every cell expressing a gene rather than only those already using multiple isoforms. Each line joins one sample’s mean entropy in its *CDKN2A*-negative and p16^INK4a^-positive arms; the slope of each line represents the effect for that sample. **(D)** Per-gene change in isoform entropy between p16^INK4a^-positive and *CDKN2A*-negative cells against the Benjamini–Hochberg adjusted p-value, on library-size-matched cells and with entropy averaged over every expressing cell. Restricted to genes whose effect direction reproduces across donors, assessed independently on the matched cells. **(E)** Per-cell isoform diversity for TES, pooled across all cell types, shown by donor and *CDKN2A* status, with the pooled comparison at right. **(F)** The percentage of cells expressing TES gene that detect 1, 2, 3, or at least 4 of its isoforms. Entropy is averaged over every expressing cell rather than only those already using multiple isoforms, so the two panels describe the same set of cells.

Non-canonical junctions were overwhelmingly unannotated and rare (**Supp. Fig. 5A–B**), consistent with long-standing estimates that non-canonical dinucleotides account for under 1% of mammalian splice junctions and are frequently attributable to sequence artifacts^35^. By contrast, novel canonical junctions showed substantially greater recurrence (**Supp. Fig. 5A–B**). Of 474,899 canonical junctions, 228,226 (48.1%) were unannotated, and 89.2% of these were detected in at least two samples, with a median of nine samples per junction. Novel splice sites clustered near their annotated counterparts, consistent with recurrent alternative splice-site usage rather than predominantly stochastic alignment errors (**Supp. Fig. 5B–C**). Although novel junctions represented 57.9% of distinct junction coordinates, they accounted for only 5.7% of all junction observations, indicating that novel splicing is widespread but generally represents only a small fraction of junction observations. The contribution of novel splicing also varied across tissues, from approximately 4.5% of junction observations in prostate to 9% in testis (**Supp. Fig. 5D**).

Transcript boundaries were similarly diverse. Observed start and end positions were offset from their reference counterparts in both directions, with tighter agreement at 3’ than at 5’ ends (**Supp. Fig. 5E–F**). Genes used multiple transcriptional start sites, with an average of 8.9 distinct starts per gene at 50 bp resolution, and Cap Analysis of Gene Expression (CAGE) and polyadenylation signals independently supported a substantial fraction of observed transcript boundaries (**Supp. Fig. 5E–I**). Notably, novel isoforms showed strong boundary support, with CAGE peaks supporting 54.5% of their start sites and polyadenylation motifs present upstream of 79.8% of their cleavage sites. Thus, novel transcript boundaries were not less well supported than annotated boundaries, arguing against degradation or mispriming as the primary source of transcript novelty.

We next sought independent evidence that novel junctions represented genuine transcripts rather than long-read artifacts. Across the four donors with multi-organ contributions, matched short-read libraries were available for 48 samples, and validation used the long-read junctions from these 48 samples only. These samples nonetheless contained 223,567 of the 228,226 distinct novel canonical junctions in the atlas (98.0%), making the validated set nearly comprehensive. Matched short-read data confirmed 154,843 of 223,567 distinct novel canonical junctions (69.3%) in at least one sample, with validation increasing with long-read support (**Supp. Fig. 6A–C**). At single-cell resolution, short reads detected a novel junction in 10.4% of cells expressing the corresponding long-read isoform, compared with only 1.3% of cells expressing the gene without that isoform, representing a ∼7.8-fold enrichment (**Supp. Fig. 6D**). Independent public RNA-seq datasets provided further support: 91.4% of novel canonical junctions were recovered in the Sequence Read Archive (SRA) compilation and 84.4% in Genotype-Tissue Expression (GTEx) (**Supp. Fig. 6E–F**). Together with the matched short-read validation, these observations across independent cohorts and sequencing platforms support the authenticity and reproducibility of the novel canonical splicing landscape.

### Isoform usage defines cellular identity beyond gene expression

An isoform’s abundance in a cell is the product of its gene expression abundance and the cell’s choice of splicing. Thus, analyzing isoform expression measures both simultaneously. We therefore asked whether isoform usage carries information beyond gene abundance. Among the most variable isoform features, isoform fractions were largely independent of gene expression, with a median Spearman correlation of −0.08 between isoform fraction and total gene abundance; 89.4% of isoforms had an absolute correlation below 0.2 (**Supp. Fig. 7A**). The weak dependence at low counts largely reflected quantization of isoform fractions rather than a biological relationship, with dominant and minor isoforms showing only small opposite correlations with gene abundance (median ρ = −0.09 and 0.06, respectively; **Supp. Fig. 7B**). We next embedded cells using isoform fractions alone, excluding gene-level expression. Cells segregated by cell type (**Fig. 3A**), and when both representations were restricted to the same set of genes, isoform fractions achieved the same mean k-nearest-neighbour cell-type purity as gene expression (0.73 for each; **Supp. Fig. 7C**). However, only half of a given cell’s neighbours were shared between the two representations (median overlap 0.50; **Supp. Fig. 7D**). Thus, the two representations achieved comparable cell-type purity despite exhibiting substantially different local structure, indicating that comparable cell-type classification arose from distinct molecular features. Thus, isoform usage captures cellular identity beyond gene expression.

To identify the genes underlying this structure, we tested isoform usage across broad cell classes using a Dirichlet-multinomial model, which estimates changes in isoform proportions independently of total gene expression (**Methods**). Across the 30 broad cell classes represented by cell types with at least 100 cells, we found 150,962 significant isoform-cell class pairs spanning 53,659 isoforms and 16,057 genes (absolute isoform-fraction change > 0.1, and a per-gene adjusted p < 0.01 broadcast to that gene’s isoforms). Nearly half of these isoforms (48.3%) were significantly differentially used in a single cell class, whereas the genes carrying them were shared across a median of four classes (**Supp. Fig. 7E**), indicating that cell types more often redistribute isoform usage among a common set of genes than alter overall gene expression.

Isoform usage can change without altering total gene expression. To distinguish these effects, we tested the same 250,159 gene–cell-class pairs for differential isoform usage (DIU) and differential expression (DE), using an absolute isoform-fraction change > 0.1 with adjusted p < 0.01 for DIU and | log□ fold-change| > 0.5 with adjusted p < 0.05 for DE. Of 81,939 significant DIU pairs, 28,469 (34.7%) occurred without significant DE (**Fig. 3B; Supp. Fig. 7F**), including 12,138 (14.8% of DIU events) with no detectable expression change. These were not simply near-threshold DE events, as their median absolute log□ fold-change was 0.10, compared with 0.63 across all tested pairs, indicating that most fell well below the 0.5 DE threshold (**Supp. Fig. 7G–H**). DIU and DE therefore capture partly overlapping but distinct regulatory changes (Jaccard index = 0.31), with only a weak correlation between isoform-usage and expression effects across pairs showing DIU (Spearman ρ = 0.23; **Supp. Fig. 7G**). Conversely, 63.3% of DE pairs show no accompanying isoform shift. The strongest usage changes also differ across cell classes (**Supp. Fig. 8A**). As examples, for S100A16 and FBXL13 expression among the cells that express each gene was comparable across cell classes, whereas the isoform composition of that expression differed markedly between them (**Fig. 3C–D**). Thus, isoform usage represents a partly overlapping but non-redundant layer of cellular regulation that cannot be inferred from gene expression alone.

We next asked whether genes differ not only in which isoforms they express, but also in how broadly their expression is distributed across isoforms. We quantified isoform diversity using Rényi collision entropy (α = 2) of each gene’s isoform fractions within each cell type, which downweights lowly expressed isoforms and emphasizes isoforms that meaningfully contribute to gene expression (**Methods**). Diversity ranged from 0 to 5.56 bits across gene–cell-type pairs and was strongly right-skewed, with most genes dominated by a small number of isoforms and a minority showing sustained multi-isoform usage (**Supp. Fig. 9A**). Male germ cells showed the highest diversity, with spermatocytes and spermatids ranking first and second among 128 cell types (mean Hα = 1.37 and 1.35, respectively, versus 1.18 median), and testis ranking highest among tissues (1.38 versus 1.33 for the next tissue; **Fig. 3E; Supp. Fig. 9B**), as previously described in the mammalian testis^36^. Genes also differed in how diversity varied across cell types, ranging from consistently high or low diversity to cell-type-restricted diversification (**Fig. 3F**); these patterns were reproducible across donor subsets (Spearman ρ = 0.73; **Supp. Fig. 9C**). Highly variable-diversity genes were enriched for novel combinations of annotated splice sites, whereas low, stable-diversity genes were enriched for isoforms containing unannotated splice sites, despite similar overall fractions of novel isoforms (**Fig. 3G; Supp. Fig. 9D**), and high-diversity genes showed greater within-gene variation in coding-sequence length (**Supp. Fig. 9E**). Representative genes illustrated these distinct patterns of isoform usage across cell types (**Supp. Fig. 9F**). Estimates were robust to alternative entropy formulations, with cell-level and cell-type-mean entropy nearly identical and Shannon entropy closely concordant, whereas Tau showed more moderate agreement (**Supp. Fig. 10A–C**). Isoform diversity therefore appears to be a reproducible feature of cellular identity, and not an artifact of the entropy measure used.

We next partitioned isoform diversity and composition across 640 cell type–sample profiles into cell-type, tissue and donor effects using degrees-of-freedom-corrected partial ω^2^ (**Methods**). Isoform diversity was more strongly associated with cell type than tissue (mean ω^2^ = 0.062 versus 0.047), whereas isoform composition showed comparable contributions from cell type and tissue after accounting for factor complexity (**Fig. 3H; Supp. Fig. 10D–F**); donor effects on diversity were smaller. Cell-type-by-tissue interactions explained a substantial fraction of residual variation, indicating that isoform regulation is not fully additive across these factors, while within-sample analyses confirmed larger cell-type effects when tissue and donor were held constant (**Supp. Fig. 10D–E; Methods**). The relative contributions also depended on annotation resolution: coarsening from 144 fine cell types to 34 broad classes shifted variance toward tissue, consistent with many fine cell types being tissue-restricted (**Supp. Fig. 10G**). Thus, isoform diversity is particularly associated with cellular identity, whereas isoform composition reflects both cell type and tissue context, with substantial context-dependence in isoform regulation.

### Isoform-level remodeling in cells expressing senescence-associated transcripts

*CDKN2A* encodes two major splice products, p16^INK4a^ and p14^ARF^, which engage distinct effector pathways. Isoform-level resolution is therefore essential for distinguishing a senescence-associated p16^INK4a^ program from a p53-mediated stress response. Because *CDKN2A* transcripts are expressed at low abundance, we enriched *CDKN2A* molecules from 10x cDNA libraries generated from four multi-tissue donors and performed deep long-read sequencing (**Methods**). This targeted approach increased *CDKN2A* molecule recovery 3.4–8.7-fold relative to matched whole-cell libraries (**Supp. Fig. 11A**), enabling sensitive assignment of *CDKN2A* status from full-length transcripts (**Methods**; **Fig. 4A–B**; **Supp. Fig. 11B–C**), while maintaining the link to cell bar code for further transcriptome analysis.

We first asked whether individual cells favor one *CDKN2A* isoform. Among cells with an identifiable p16^INK4a^ or p14^ARF^ transcript, the majority expressed both isoforms rather than either isoform alone (2,565 of 3,157 cells, 81.2%). The two isoforms co-occurred more frequently than expected by chance, with a per-sample enrichment of 28.6-fold over an empirical null (95% CI 16.0–51.3; p = 4.3 × 10⁻¹□), or 21.2-fold with donor as the unit of replication (95% CI 12.5–36.1, four donors; **Fig. 4B, Supp. Fig. 11D**)^28^. This is consistent with the two promoters being embedded in a single Polycomb-repressed chromatin domain that is derepressed as a unit^37^, and extends that coordination from the population level to individual cells. Among cells in which a first exon could be assigned, 99.3% (3,134 of 3,157) expressed p16^INK4a^, indicating that detection of p14^ARF^ was nested almost entirely within p16^INK4a^-positive cells. Together, these results indicate that individual cells generally co-express both *CDKN2A* isoforms rather than selectively expressing one.

We next asked whether the p16^INK4a^-associated senescence program is accompanied by remodeling of transcript isoform architecture. We previously identified *CDKN2A*+ MKI67-cells with multiple hallmarks of senescence and heterogeneous senescence phenotypes using short-read RNA sequencing^12^. Here, we defined senescence-associated cells by detectable p16^INK4a^ transcripts, including p16^INK4a^-only and p16^INK4a^/p14^ARF^ double-positive cells (n = 3,134), and compared them with cells of the same type lacking detectable *CDKN2A* transcripts (n = 151,048). Cells expressing p14^ARF^ alone (n = 23) or other *CDKN2A* transcripts without p16^INK4a^ (n = 912) were excluded, ensuring that all comparisons were defined by p16^INK4a^ status.

Individual p16^INK4a^-positive cells expressed marginally more transcript isoforms than their *CDKN2A*-negative counterparts. To avoid the strong dependence of aggregate isoform counts on the number of cells sampled and on sequencing depth, we compared groups at matched cell number and matched depth. At matched cell number, the two groups differed only marginally in the total number of distinct isoforms detected (1.02-fold), and this small excess was confined to previously unannotated structures (1.04 for NNC isoforms against 1.01 or below for FSM, ISM, and NIC isoforms; **Supp. Fig. 11E**). The difference was clearer per cell, a measure that does not depend on how many cells each group contributes: p16^INK4a^-positive cells detected 158 more isoforms per cell (95% CI 64–252; p = 1.7 × 10^−3^ with sample as the unit of replication; medians 2,372 versus 2,195), a difference present in 31 of the 36 estimable samples.

We quantified per-cell isoform diversity as the mean Shannon entropy of isoform usage, averaged over every gene the cell expresses (**Methods**). p16^INK4a^-positive cells showed higher per-cell entropy than *CDKN2A*-negative cells (+0.014 bits, 95% CI 0.008–0.020; p = 4.3 × 10^−5^ with sample as the unit of replication), with the effect present in 30 of 36 samples (**Fig. 4C**), mirroring the entropy gain reported in cancer^38^. The increase was reproduced in 19 of the 22 broad cell classes estimable on the matched cells (of 34 classes represented in the atlas; **Supp. Fig. 11F**), arguing against an explanation based solely on enrichment of cell types with intrinsically high isoform complexity. Whereas aggregate isoform counts capture differences in transcript repertoires across cells, per-cell entropy indicates that individual p16^INK4a^-positive cells themselves co-express a broader range of isoforms.

Gene-level analyses likewise revealed a global shift toward increased isoform diversity in the p16^INK4a^ program (**Fig. 4D**). Because diversity could be assessed only for genes with at least two detected isoforms, and because p16^INK4a^-positive cells were less abundant than their comparators, the expected fraction of genes with increased diversity under the null was 48.7%, estimated by permuting p16^INK4a^ status within samples 1,000 times. In the observed data, 59.3% of the 14,236 testable genes showed increased diversity, a 10.6-percentage-point excess corresponding to 3.3 null standard deviations. This shift was reproducible across individuals, with 4,458 genes changing in the same direction in at least three of four donors. Although significant effects were often restricted to individual cell classes (**Supp. Fig. 12A**), this apparent specificity was strongly influenced by sampling: power analysis indicated that 73.5% of non-significant class–gene comparisons were consistent with insufficient power rather than absence of an effect.

Increased entropy reflected broader within-cell co-detection of isoforms. For example, *TES*, which had 10 detected isoforms across the atlas, showed co-detection of two or more isoforms in 40.9% of p16^INK4a^-positive cells expressing *TES*, compared with 18.1% of *CDKN2A*-negative cells expressing *TES* (**Fig. 4E–4F**). Thus, increased isoform diversity arose primarily through coordinated expression of multiple isoforms within individual cells rather than redistribution among a fixed set of isoforms. The same pattern was evident in cell-class-specific examples, including FN1 in fibroblasts, for which the comparison was estimable in all four donors (**Supp. Fig. 12B**) and whose splicing was previously shown to vary between individual cells and to shift during cellular senescence^39^, and CTSL in myeloid leukocytes, which showed one of the largest entropy increases in that class and increased in all three donors in which it was estimable (**Supp. Fig. 12C**)^40^.

In addition to increasing overall isoform diversity, the p16^INK4a^ program was associated with directed changes in isoform usage. Differential isoform usage (DIU) analysis within each broad cell class identified 4,006 genes with significant isoform switches (**Supp. Fig. 12D**). Most genes showed significant differential isoform usage in only one class (79.3%), and the number of detected switches strongly tracked the number of p16^INK4a^-positive cells in each class (Spearman ρ = 0.91). However, this apparent cell-class specificity was substantially reduced when effects were evaluated on a common scale across classes. For 69.0% of non-significant class-gene pairs, confidence intervals remained compatible with the effect estimated in the discovery class, and the median gene had an informative readout in three classes. Moreover, 82.7% of genes reached at least half of their discovery-class effect in at least one additional class. On this common scale the fraction of genes apparently restricted to a single class fell from 79.3% to 47.3%. Equalizing cell numbers across classes independently reduced the correlation between switch detection and cell number from 0.91 to 0.27 at 50 cells per class, although at equal n, far fewer genes remained significant and the single-class fraction rises, as expected when power falls. These results indicate that the apparent restriction of p16INK4a-associated isoform switching to single cell classes is substantially inflated by unequal detection power across cellular contexts, although roughly half of switch genes remain class-restricted once effects are placed on a common scale.

Together, these analyses reveal a coordinated remodeling of transcript isoform architecture in p16^INK4a^-positive cells. Rather than selectively activating a single *CDKN2A* isoform, individual cells generally co-express p16^INK4a^ and p14^ARF^, while the broader p16^INK4a^-associated program is characterized by increased within-cell isoform diversity, expansion of previously unannotated transcript structures, and widespread changes in isoform usage across cell types.

## Discussion

This cross-tissue single-cell long-read isoform atlas provides an isoform-resolved counterpart to gene-level references across human tissues. Harmonization of transcript structures across samples enables direct comparison of isoform usage across cell types, tissues and individuals. A substantial fraction of detected transcript structures is absent from current annotations, yet these previously unannotated isoforms recur across independent samples and are supported by orthogonal evidence, demonstrating that full-length single-cell sequencing can expand reference transcriptomes with biologically reproducible transcript structures.

Isoform usage also defines a distinct axis of cellular identity. Cell types showed widespread differences in transcript structure, including among genes with similar overall expression. Isoform repertoire was more strongly associated with cell type than tissue of residence, whereas individual isoform usage reflected both lineage and tissue context. These patterns were reproducible across donors, establishing isoform usage as an independent and structured layer of cellular identity.

The biological utility of isoform-level resolution is illustrated by *CDKN2A*, where full-length sequencing revealed frequent co-expression of p16^INK4a^ and p14^ARF^ within individual cells and exposed cell-type-specific isoform remodeling associated with the p16^INK4a^ senescence program. In most somatic tissues, short-read *CDKN2A* detection provides a useful proxy for p16^INK4a^ expression, but it cannot resolve the distinct transcript products generated from this locus, a distinction that becomes critical in tissues such as testis, where p19^ARF^ expression in mice is confined to spermatogonia and extinguished in spermatocytes^41^. More broadly, resolving transcript structures at single-cell resolution provides access to coding potential, transcript stability, and regulatory features that gene-level measurements cannot distinguish, establishing a framework for studying isoform regulation across human cell states, tissues and disease contexts.

## Data and code availability

All data supporting the findings of this study are available within the article and its supplementary information files or can be obtained from the corresponding author upon reasonable request. The raw sequencing data will be deposited in a public AWS S3 bucket upon publication. To protect donor genetic privacy, access to the raw sequencing reads will require a data transfer agreement.

All gene counts, isoform counts, and single-cell metadata generated from the long-read sequencing data have been made publicly available through a Figshare repository (https://doi.org/10.6084/m9.figshare.33411454). The analysis pipeline, along with all scripts and notebooks required to reproduce the analyses presented in this study, is publicly available in a GitHub repository (https://github.com/madhavmantri/tabula_longread).

## Acknowledgements

This project was supported in part by grant nos. 2019-203354, 2020-224249, 2021-237288, 2021-006486, 2022-316725 from the Chan Zuckerberg Initiative DAF, an advised fund of Silicon Valley Community Foundation, and by support from the Chan Zuckerberg Biohub San Francisco. We thank Donor Network West for procuring the organs and tissues from the donors for this project. We are also grateful to the UCSF Liver Center (funded by NIH P30DK026743) for assistance with the liver cell isolations, and to B. Tojo for the original artwork in Figure 1. We thank members of the Quake lab for many valuable discussions. We also thank Pacific Biosciences and Signios Biosciences for their support with long-read sequencing.

## Author Contributions

S.R.Q. supervised the study. M.M., A.M.D., N.F.N., R.C.J. and L.T. performed the PacBio Kinnex long-read sequencing experiments. The Tabula Sapiens Consortium provided the scRNA-seq cDNA used for the PacBio Kinnex experiments. M.M., J.L., L.T., J.H.Y.K. and A.Z. analysed the data. M.M. and S.R.Q. wrote the manuscript with input from all authors. All authors reviewed and approved the final manuscript.

## Declaration of interests

The authors declare no competing interests.

## Methods

### Organ and tissue procurement

To maintain consistency in the overall Tabula Sapiens dataset, organ and tissue procurement followed the same procedure used in the first phase of the project. Donated organs and tissues were procured at various hospital locations in the Northern California region through collaboration with a not-for-profit organization, Donor Network West (DNW, San Ramon, CA, USA). DNW is a federally mandated organ procurement organization (OPO) for Northern California. Recovery of non-transplantable organs and tissues was considered for research studies only after obtaining records of first-person authorization (i.e., donor’s consent during his/her DMV registrations) and/or consent from the family members of the donor. However, the pancreas from donor TSP9 was provided by Stanford University Hospital under appropriate regulatory procedures. The research protocol was approved by the DNW’s internal ethics committee (Research project STAN-19-104) and the medical advisory board, as well as by the Institutional Review Board at Stanford University which determined that the Tabula Sapiens projects do not meet the definition of human subject research as defined in federal regulations 45 CFR 46.102 or 21 CFR 50.3.

### Preparation of single-cell suspensions

Tissues were obtained from 12 Tabula Sapiens donors and processed as previously described in the Tabula Sapiens atlas^10,12^. Single-cell suspensions were prepared from freshly dissociated tissue as described in Tabula Sapiens. At least two donors were sampled per tissue for long-read RNA sequencing, with one male and one female donor for every non-reproductive tissue except ear and pancreas, which were sampled from male donors only.

### Long-read library preparation and sequencing

Long-read libraries were prepared from 60 distinct samples spanning 26 tissues. A 15 ng aliquot of the barcoded, full-length cDNA generated by the 10x Genomics 3’ RNA-seq reaction was used for long-read library preparation, such that long-read RNA sequencing was performed on material derived from the same dissociated cells as the short-read datasets. TSO-TSO (template switching oligo) artifacts were depleted from the full-length cDNA pool. The cDNA fragments were then concatenated into 16-segment concatemer molecules, followed by SMRTbell adapter ligation using the PacBio Kinnex single-cell RNA kit (formerly MAS-Seq; #103-072-200) according to the manufacturer’s instructions. Libraries were sequenced on a PacBio Revio instrument to generate circular-consensus (HiFi) reads. Sequencing was performed by Pacific Biosciences and Signios Biosciences.

### Per-sample Iso-Seq processing

Raw HiFi reads were processed through a unified Snakemake^42^ pipeline (v7.32.4). Concatemeric HiFi reads were first split into their constituent cDNA segments with *skera split* (pbskera v1.4.0) using the MAS-Seq v1 16-mer adapter FASTA, and the segmented BAM files from all input movies of a sample were merged. The 5’ and 3’ primers were then removed, and reads oriented with *lima --isoseq* (lima v2.13.0) against the MAS-Seq 16-mer primer FASTA. Cell barcodes and UMIs were extracted with *isoseq tag* (isoseq v26.2.0) under the *T-12U-16B* read design, corresponding to a 12 bp UMI followed by a 16 bp cell barcode, after which reads were refined to full-length non-concatemer transcripts with *isoseq refine --require-polya*. Cell barcodes were corrected against the 10x 3M-february-2018 whitelist, reverse-complemented to match the orientation in which PacBio stores barcodes, using *isoseq correct --method percentile --percentile* 90, and real cells were called at the same percentile with *isoseq bcstats*, where the UMI cutoff is defined as ten times the UMI count at the top 10% barcode rank. Corrected reads were sorted by the corrected cell-barcode tag (CB) with *samtools sort -t CB* (samtools^43^ v1.22.1). Five libraries (T4, T13, T21, T22, and T52) carried more than ten real-cell barcodes above 200,000 reads and were capped at 200,000 reads per cell barcode before deduplication for groupdedup command efficiency. Reads sharing a cell barcode, UMI and insert structure were then collapsed into single molecules with *isoseq groupdedup*, deduplicated molecules aligned to the GRCh38.p13 primary assembly with *pbmm2 align --preset ISOSEQ --sort* (pbmm2 v26.1.99) which uses minimap2 aligner^44^, and aligned molecules collapsed into per-sample transcript models with *isoseq collapse --do-not-collapse-extra-5exons --max-5p-diff1000 --max-3p-diff1000.* The reference genome was the GENCODE GRCh38.p13 primary assembly and the GENCODE v41 reference annotation^45^. The same FASTA and GTF were used for every alignment, classification and short-read index in this study, so that long-read and short-read coordinates are directly comparable.

### Cross-sample isoform identifier harmonization

Independent per-sample collapse generates sample-specific isoform identifiers, which cannot be compared across tissues or donors. Identifiers were therefore harmonized by a global re-collapse. Each sample’s collapsed transcript FASTA was prefixed with its sample of origin and the prefixed files were pooled across all samples. The pooled sequences were re-aligned to GRCh38.p13 with *pbmm2 align --preset ISOSEQ* and re-collapsed globally with *isoseq collapse* under the same parameters used per sample. The resulting global identifiers were written back onto each sample’s original collapse outputs, so that each transcript structure retained a single identity atlas-wide while preserving per-sample molecule assignments. This rewrite step was verified to conserve counts. The relabelled models were then passed through the same *pigeon* (pbpigeon v26.2.0) sort, classify and filter workflow and Seurat matrix construction described below.

Because the Seurat matrix is rebuilt from the filtered classification rather than copied, global merging can flip individual filter decisions in either direction. Measured across all 60 samples, the recollapsed feature totals differed from the per-sample totals by −0.027% overall, with 30 samples up and 30 down, a mean absolute per-sample change of 0.39% and a maximum of 4.03%. The global re-collapse merges structures across samples by coordinate. Recurrence statistics, defined as the fraction of isoforms detected in two or more samples, are computed after this step and after the atlas-wide abundance filter described below, with known isoforms passing through the identical procedure as the matched control.

### Transcript classification and functional annotation

Harmonized transcript models were classified against GENCODE v41 with pigeon classify, supplying per-transcript full-length molecule counts, the refTSS^46^ v3.1 hg38 CAGE-peak set and the combined human and mouse poly(A) motif list, which encodes the canonical AAUAAA hexamer and its common variants^47^, and were then filtered with pigeon filter. Functional annotation was transferred onto the filtered models with IsoAnnotLite^48^ (v2.7.4). All coding, ORF, CDS, and predicted-NMD annotations were obtained from SQANTI3 package^49^ v6.0.1, run as a locally patched build, with the CAGE-peak and poly(A)-motif data files taken from the SQANTI3 v5.5.4 data bundle. SQANTI3 quality control was run with *sqanti3_qc.py --include_ORF*, supplying the refTSS v3.1 hg38 CAGE peaks *(--CAGE_peak)*, the human and mouse poly(A) motif list *(--polyA_motif_list)* and a BED file of annotated polyadenylation sites built from the GENCODE v41 *polyA_site* features *(--polyA_peak)*, with full-length counts given as deduplicated molecule counts only. Rules-based filtering was applied with *sqanti3_filter.py rules*.

Structural categories follow the SQANTI3 definitions used throughout, namely full-splice match (FSM), incomplete-splice match (ISM), novel-in-catalog (NIC), novel-not-in-catalog (NNC), fusion, moreJunctions, genic, antisense and intergenic. Two conventions should be noted when reading the resulting numbers. First, *reference_match* requires a transcript’s 5’ and 3’ ends to fall within SQANTI3’s default 50 bp tolerance of the reference transcript, not to be coordinate-identical, so the figures described in Results as matched at both ends are matches within that tolerance. Second, transcript coding-sequence lengths were taken from the SQANTI3 *CDS_length* field. The SQANTI3 *ORF_length* field reports the genomic span of the CDS rather than the summed length of its exonic blocks and was not used.

### Count matrix construction and feature levels

Cell-by-feature matrices were generated with pigeon make-seurat, with novel genes and ribosomal and mitochondrial genes excluded. A second matrix was generated with --keep-pbids, in which every feature carries the PacBio identifier that links it to the collapse group file. Four nested feature levels were produced from these matrices: genes, pbids, ensemblids, and ensemblids_annotatedonly. These were converted to AnnData objects for each donor, retaining cells with at least 500 counts. The gene level holds one feature per gene. The pbid level holds one feature per distinct collapsed transcript structure and includes both annotated and novel structures. The ensemblid level is formed by summing pbid counts within each transcript identifier, so that structures assigned to the same GENCODE transcript are pooled while structures matching no annotated transcript retain their PacBio identifier. The annotated-ensemblid level is the subset of the ensemblid level carrying an Ensembl transcript identifier, and therefore excludes novel structures. Per-donor objects were then concatenated into all-sample objects, with .var annotation rebuilt from the union of donors so that features contributed by any donor retain their gene and transcript annotation.

The gene level underlies cell filtering, the atlas cell counts and the differential expression analysis. Three isoform levels are used for the analyses reported here. The pbid level comprises all 854,410 distinct transcript structures across 37,805 genes and was used to establish the novel isoform and splicing landscape and to compare the structural and predicted functional properties of isoforms. The ensemblid level comprises 489,842 features across 38,590 genes and was used for the usage analyses that require a stable isoform composition per gene, namely isoform diversity and the isoforms that encode cell identity. The annotated-ensemblid level comprises 145,051 features across 33,676 genes and was used for the senescence analyses, namely isoform entropy and differential isoform usage between senescent and non-senescent cells. The usage and senescence analyses are computed on the isoform-fraction restriction of their level, which retains only genes with at least 2 isoforms and leaves 478,192 features over 26,940 genes at the ensemblid level and 130,362 features over 18,987 genes at the annotated-ensemblid level. All three are referred to as isoforms in Results, and every quoted total identifies which population it describes.

### Quality control and cell filtering

Filtering was applied in a fixed order, with all decisions taken on the gene-level matrix and inherited by the finer levels, so that the four levels retained the same set of cells. An ambient and index-hopping filter was applied first, removing cells whose 10x barcode collided across two long-read libraries of the same donor, and this was done on raw cells at the gene level, before any other quality-control step. Cells with more than 1,000 total UMIs on the full gene matrix were then retained, a threshold applied only at the gene level and on the true library size, with the pbids, ensemblids, and ensemblids_annotatedonly levels inheriting the surviving cell set and applying no UMI cut of their own. Among the retained cells, genes detected in at least 5 cells were kept, and every isoform belonging to a removed gene was dropped at the finer levels. Individual features at the pbid, ensemblid and annotated-ensemblid levels were additionally required to be detected in at least 5 cells and to carry at least 10 total counts across the atlas. This threshold defines the robust transcript structures used throughout, namely at least 10 unique molecules across at least 5 cells. Cell-type labels were then attached as described below and cells with a missing or unknown label were removed at the gene level. After filtering, 203,311 cells were retained.

### Cell type annotation

Cells were annotated by consensus prediction with popV^50^ (v0.6.0), using the tissue-specific Tabula Sapiens pretrained models in fast prediction mode, run independently per tissue on the gene-level matrix. popV labels were mapped through a manually curated table onto Cell Ontology^51^ terms, giving the fine cell type (*cell_ontology_class*), and each fine type was then assigned to one of the broad cell classes used by Tabula Sapiens (*broad_cell_class*). In the filtered atlas, this yielded 144 fine cell types grouped into 34 broad cell classes. Cell-type labels were incorporated into the preprocessed objects and are not re-derived by downstream notebooks.

### Isoform fractions

For every gene, the isoform fraction of isoform i in cell c was defined as the count of i in c divided by the total count of that gene’s isoforms in c, with all fractions set to zero where the gene is not detected. Fractions were stored as a sparse layer alongside the raw counts. For the usage analyses, this layer was restricted to genes retaining at least 2 isoforms.

### Matching long-read and short-read cells

PacBio cell barcodes are stored in reverse-complement orientation relative to the short-read barcodes of the same 10x run. Cells were matched across platforms by reverse-complementing each PacBio barcode and pairing it with the short-read barcode of the same 10x sample, keyed on short-read sample identifier and barcode jointly. For the platform-concordance analysis, the comparison was restricted to cells recovered in both assays and to samples with a matched short-read library. Counts were summed across shared barcodes within each donor, normalized to counts per 10,000 and log1p-transformed, and Pearson and Spearman correlations were computed across genes detected in either platform, with donor-level pseudobulks requiring at least 20 shared cells. Technical reproducibility was assessed identically on the two libraries prepared from two 10x samples of the same tissue (T29 and T30).

### Splice junctions

Splice junctions were taken from the SQANTI3 junction tables of the robust isoform set. Junctions were counted both as distinct genomic coordinates and as observations weighted by the isoforms carrying them, and were split by donor and acceptor motif into canonical (GT-AG, GC-AG, AT-AC) and non-canonical classes, and by annotation status into known and novel. All recurrence statistics are reported per motif class. Novel 5’ and 3’ splice sites were enumerated separately and the distance from each to its nearest annotated counterpart was computed.

### Transcript boundaries

Observed transcription start and termination sites were compared with their reference counterparts for full-splice-match and incomplete-splice-match transcripts, using the SQANTI3 *diff_to_TSS and diff_to_TTS* fields. Independent boundary support was taken from the SQANTI3 annotations described above, in the form of overlap of a transcript’s 5’ end with a refTSS v3.1 CAGE peak, presence of a poly(A) motif upstream of the cleavage site, and overlap of the 3’ end with an annotated GENCODE v41 polyadenylation site.

Distinct transcription start sites were counted per gene at 50 bp resolution, by binning observed start coordinates to 50 bp and counting distinct bins per gene. The per-gene average is a lower bound on promoter diversity rather than an absolute tally, since the binning merges start sites closer together than 50 bp. The average is therefore reported both across all genes carrying a start-site annotation and across well-sampled genes, defined as genes with at least ten detected transcript structures.

### Per-cell functional metrics by cell type and tissue

For each cell, the fraction of its detected molecules assigned to isoforms carrying each functional property was computed from the pbid-level matrix joined to the SQANTI3 classification. The properties considered were novel splicing (NIC or NNC), predicted coding potential, predicted NMD, retained intron, and an exon-skipping proxy defined as an isoform having fewer exons than its associated reference transcript. Cell types and tissues represented by fewer than 20 cells were dropped from the summaries. The exon-skipping metric is a proxy and should be interpreted as an upper bound on skipping, since an ISM fragment of a longer transcript also satisfies it.

### Short-read validation of novel splice junctions

For 48 of the 58 samples with a matched short-read library (restricted to the four multi-organ donors TSP21, TSP25, TSP27 and TSP33), reads were aligned with STARsolo^52^ (STAR v2.7.10b aligner^53^) against an index built from the same GRCh38.p13 primary assembly and GENCODE v41 annotation used for the long-read pipeline, with *--sjdbOverhang* 100. STARsolo was run in *CB_UMI_Simple* mode with the 10x 3’ v3.1 geometry, giving a 16 bp cell barcode from position 1 and a 12 bp UMI from position 17, against the forward-orientation 3M-february-2018 whitelist, emitting both gene and splice-junction features *(--soloFeatures Gene SJ*). *CellRanger-emulating* options were used throughout *(--soloCBmatchWLtype1MM_multi_Nbase_pseudocounts, --soloUMIdedup 1MM_CR, --soloUMIfiltering MultiGeneUMI_CR, --clipAdapterType CellRanger4, --outFilterScoreMin 30)*.

The target set comprised the novel canonical junctions of isoforms present in the preprocessed pbid-level object, so that the tested junctions are exactly those that survive the atlas-wide robustness filter. Each distinct junction was tested once in every sample in which it was detected in the long-read data, and counted as validated if a junction with identical chromosome, start and end coordinates was present in that sample’s STARsolo SJ output. Two controls were run using the same procedure, a positive control of known canonical junctions from the same samples, and a negative control of coordinate-shifted decoys constructed by displacing both ends of every target junction by +137 bp. Validation rate was additionally tabulated against long-read support, binning each distinct junction by the total number of molecules supporting it summed over all isoforms that carry it, and against relative position within the isoform’s genomic span. The short-read libraries are 3’-biased, so the positional analysis is interpreted through the gap between novel and known junctions at matched position, with the known curve as the estimate of detection power.

Agreement at single-cell resolution was assessed separately. For each isoform and junction pair, the short-read detection rate of the junction was compared between cells expressing that isoform in the long-read data (LR+) and cells expressing the same gene without that isoform (LR−), across barcodes matched between platforms as described above. Pairs were required to have at least 5 LR+ cells, and the two rates were compared by a paired Wilcoxon signed-rank test over pairs pooled across the 48 samples. The unit is the isoform and junction pair.

### External validation against public compendia

The atlas-level set of distinct novel canonical junctions was queried against Snaptron^54^, in both the srav3h compilation^55^ (SRA human) and gtexv2 database^56^ (GTEx) compilations. The same +137 bp coordinate-shifted decoy set was queried in parallel, and the denominator restricted to junctions falling inside successfully fetched query windows so that partial coverage does not inflate the estimated recovery rate. Recovery was reported both as presence in at least one library and as a function of the minimum number of supporting Snaptron samples required to count a junction as found.

### Isoform usage versus gene abundance

For each isoform, the Spearman correlation between its isoform fraction and its own gene’s total abundance was computed across the cells in which that gene was detected. Genes were required to be detected in at least 50 cells, isoforms to be detected in at least 50 of those cells, and up to 5,000 cells were sampled per gene to limit computational cost. The analysis was run on the isoform features used for the embedding, and the analysis is therefore restricted to those features rather than to isoforms as a class. Isoforms were additionally split at a mean fraction of 0.5 into minor and dominant classes, to make visible the quantisation that arises at low gene counts, where a cell holding a single molecule forces one isoform to a fraction of one and the rest to zero.

### Isoform-fraction embedding

Highly variable isoform features were selected as the 2,000 isoforms with the highest raw variance of their isoform fraction across cells, computed on the ensemblid-level object after its restriction to genes retaining at least 2 isoforms. Principal components (50, arpack solver) were computed on the isoform fraction matrix of the selected features, a neighbour graph was built with k = 30, and a UMAP embedding (umap-learn v0.5.9.post2) was computed from it.

### k-nearest-neighbour cell-type purity

Cell-type purity was computed as the fraction of each cell’s k = 30 nearest neighbours sharing its *cell_ontology_class* label, on a random subsample of 20,000 cells. Two representations were built over the same subsample, the isoform-fraction matrix restricted to the selected isoform features, and the log1p gene-total matrix restricted to the genes those isoforms come from, so that the two representations differed only in what is the quantity measured for each gene, and not in which genes were used. Each was scaled and reduced to 50 principal components before neighbour search. The overlap between each cell’s neighbour sets in the two representations was computed on the identical subsample and k.

### Differential isoform usage across cell classes

Differential isoform usage was tested with a gene-level Dirichlet-multinomial likelihood-ratio test on isoform counts^57^, fitted at the ensemblid level for the cell-identity analyses and at the annotated-ensemblid level for the senescence analyses. For each gene, a Dirichlet-multinomial concentration vector was fitted by maximum likelihood (L-BFGS-B on the log concentration) separately to the test group and the reference, and jointly to the pooled cells. The test statistic is twice the difference in log-likelihood between the separate and pooled fits and was compared with a chi-squared distribution with K degrees of freedom, K being the number of isoforms entering the fit. Genes were required to carry at least 2 isoforms and to be detected in at least 50 cells on each side of the contrast, a symmetric floor that does not bias which isoforms are visible within a tested gene. Genes with more than 30 isoforms had all but the 29 most abundant were combined into a single ‘other’ category, to keep the degrees of freedom bounded. One Benjamini–Hochberg-adjusted p-value was obtained per gene and broadcast to that gene’s isoforms, and a signed per-isoform change in mean isoform fraction was reported alongside it.

A gene-level test was chosen in preference to per-isoform testing because a per-isoform testability floor excludes an isoform lost in the test group, so only gains are recoverable; the Dirichlet-multinomial test has no per-isoform gate and recovers gains and losses together. Each contrast compared one broad cell class against all other cells, restricted to cells belonging to fine cell types with at least 100 cells, which leaves 30 broad cell classes testable. An isoform was called differentially used when its gene’s adjusted p-value was below 0.01 and its own absolute change in isoform fraction exceeded 0.1.

### Differential gene expression and the usage versus expression comparison

Gene-level differential expression was computed on the gene-level matrix normalized to 10,000 counts per cell and log1p-transformed, testing each broad cell class against all other cells with *rank_genes_groups* (scanpy^58^ v1.11.5) and correcting by Benjamini–Hochberg. A gene and cell class pair was called differentially expressed when its adjusted p-value was below 0.05 and its absolute log_2_ fold change exceeded 0.5.

The two axes were compared on an identical universe of gene and cell class pairs testable on both, so that the four-way partition into usage only, expression only, both, and neither shares a single denominator. The usage-only share depends on the gene-level fold-change gate, so the thresholds are stated wherever this quantity is quoted and the value reported in the Results corresponds to the thresholds used in the associated panel.

### Isoform diversity

Isoform diversity was quantified as the Rényi entropy of order α = 2 (collision entropy) of a gene’s isoform-usage vector, defined as H_α_ = log_2_ ( Σ_i_ p_i_^α^) / (1 − α), which for α = 2 reduces to −log_2_ Σ p_i_^2^ and places all genes on a common scale from single-isoform dominance (0 bits) to balanced multi-isoform usage. Entropy was computed per gene and cell type. For each cell type with at least 10 cells, the mean isoform-fraction vector of each gene was formed over the cells expressing that gene, and the entropy of that vector taken. Genes were required to have at least 2 isoforms and to be expressed in at least 5 cells of the cell type. The same procedure was applied per tissue for the tissue-level ranking.

Genes were then summarized by the mean and the variance of their Rényi entropy across the cell types in which they were testable, and split at the median of each axis into four quadrants. The quadrants differ in the number of isoforms per gene, and entropy is bounded above by log_2_ (number of isoforms), so they are partly isoform-count strata, and every quadrant comparison reported is therefore also shown stratified by isoform count. Reproducibility of the variance that defines these quadrants was assessed by a split-half analysis. Donors were assigned alternately by rank of cell count to two halves of six donors each, so that the halves carry comparable cell numbers. The entire per-gene and per-cell-type entropy table was then recomputed independently within each half under the identical thresholds, and the across-cell-type variance of entropy correlated between halves by Spearman correlation, overall and within each quadrant. The isoforms of each quadrant’s genes were characterised by their SQANTI3 structural subcategory mix, by the fraction of each gene’s isoforms that are novel, by predicted coding fraction and NMD rate, and by the within-gene coefficient of variation of *CDS_length* among each gene’s coding isoforms.

### Robustness of the diversity metric

Three robust analyses were performed. The scalar entropy computed on the cell-type-mean usage vector was compared with a matrix-based Rényi entropy that incorporates per-cell heterogeneity, with both averaged within a gene across the cell types in which it is expressed. Conclusions were then checked against Shannon entropy and normalized Shannon entropy computed on the same usage vectors. They were finally checked against the Tau specificity index^59^, with that comparison stratified by the number of isoforms per gene.

### Cell type versus tissue variance partition

Individual cells do not carry enough molecules per gene to estimate isoform diversity reliably, so the partition was performed on pseudobulk profiles. For every fine cell type and sample combination, the mean isoform-fraction vector of each gene was computed over the cells of that combination expressing it, requiring at least 10 expressing cells, and each profile was labelled with the tissue and donor of its sample. The unit is the cell type and sample pair rather than the cell type and tissue pair, every tissue carrying at least two samples. Genes were analysed if they carried at least 2 isoforms and at least 2 pseudobulk profiles, and were retained only where both factors varied over at least 2 levels and the residual degrees of freedom were positive. This gives 640 profiles spanning 144 fine cell types, 60 samples and 26 tissues, and retains 10,850 genes. An additive model with cell type, tissue and donor was fitted per gene using drop-first dummy coding and type-II sums of squares, with two response variables run side by side. Isoform composition was analysed by distance-based PERMANOVA on Bray–Curtis dissimilarities, the dissimilarity matrix being Gower-centred (G = −½ J D^2^ J) and factor sums of squares obtained from projection-matrix traces. Isoform diversity was analysed by two-way analysis of variance on the Rényi entropy (α = 2) of each profile. Both arms are reported as degrees-of-freedom-corrected partial ω^2^, computed as (SS_k_ − df_k_ · MS_residual_) / (SS_total_ + MS_residual_) and truncated at zero. Raw R^2^ and η^2^ are not degrees-of-freedom adjusted and favour whichever factor contributes more dummy columns. Raw values are shown in the supplement and are descriptive only.

The additive residual was further partitioned into a cell-type-by-tissue interaction and unmodelled variation by comparing the additive fit with a cell-means fit over the observed cell type and tissue cells, which requires at least one replicated cell type and tissue combination. The unmodelled term is not pure error, since each sample is nested within a single tissue and donor combination and donor-by-cell-type and three-way effects land there inseparably. Two further fits are reported as checks rather than as alternative decompositions. A two-level partition estimates cell type within sample, with sample absorbing library, tissue and donor, and estimates tissue and donor between samples on one row per sample. A resolution ladder refits the partition with the cell-type factor coarsened from 144 fine cell types to 34 broad cell classes, holding tissue fixed and rebuilding the pseudobulk profiles at each resolution, and omits the donor term. Neither is comparable with the additive estimates or with the other, being fitted on different units, gene sets or factor sets. Type-II sums of squares leave covariance shared between correlated factors unattributed, so per-factor terms do not sum to one. The diversity arm was also fitted on Shannon entropy as a concordance check. Agreement between the two models was assessed as the Spearman correlation of the cell-type-minus-tissue ω^2^ difference between them. Rényi entropy at α = 2 is the metric reported throughout, and Shannon entropy is retained only for this comparison.

### CDKN2A targeted capture

*CDKN2A* molecules were enriched from 10x cDNA generated from four multi-tissue donors (TSP21, TSP25, TSP27, and TSP33) by hybridization capture using a *CDKN2A*-targeted probe panel comprising a pooled 4 nmol synthesis of 10 Ultramer DNA oligos (IDT). Each probe contained a 5′ biotin modification (/5Biosg/) and targeted an exon across the annotated *CDKN2A* isoforms. Hybridization capture was performed using xGen™ Hybridization and Wash Reagents v3 (IDT #10028311) and xGen™ Hybridization and Wash Beads v3 (IDT #10025272), following the manufacturer’s protocol. For each capture reaction, 3 pmol of the pooled probe panel was used, corresponding to 4 µL of a 0.75 pmol/µL probe stock. 10x 3′ and 5′ primers, together with TTTTTTTTTTTTTTTTTTTTTTTTTTTTTT/3InvdT/, were included as blockers to reduce nonspecific hybridization to library adapters and poly(A) sequences. Captured material was used for library preparation and deeply sequenced on a PacBio Revio instrument using the same Kinnex chemistry as the whole-cell libraries.

Captured libraries were processed through a self-contained copy of the Iso-Seq pipeline with three parameter changes appropriate to a targeted and extremely heavy-tailed barcode distribution. Reads were capped at 10,000 per cell barcode before deduplication, real cells were called at percentile 0, and deduplication was sharded 100 ways by cell barcode using a longest-processing-time bin-packing plan and merged afterwards. The same 10,000-read cap is applied when planning the shards, since the bin-packing plan is computed from capped read counts and would otherwise not match the subsampled data.

### CDKN2A isoform assignment and cell classification

*CDKN2A* isoforms were assigned to arms of the locus by their pigeon associated_transcript, using the annotated p16^INK4a^ transcripts (ENST00000304494, ENST00000494262, ENST00000498628, ENST00000498124, ENST00000578845, ENST00000380151) and the annotated p14^ARF^ transcripts (ENST00000579755, ENST00000530628). *CDKN2A* isoforms matching neither set were labelled *other_CDKN2A*. Gene assignment was exact on the symbol *CDKN2A*, excluding *CDKN2A*-DT. Isoform matrices built with *--keep-pbids* were used for this step.

Two count definitions were retained in parallel for every cell. Raw matrix counts were taken directly from the pigeon Seurat matrices, while distinct-UMI counts were recovered by a single pass over the deduplicated FASTA, counting distinct UMI (XM) tags per cell barcode and *CDKN2A* isoform and mapping molecules to isoforms through the collapse group file. The two are not interchangeable, since isoseq groupdedup groups reads by cell barcode, UMI and insert structure and so splits a deeply sequenced UMI across several molecules. These two count measures identified the same population of cells expressing *CDKN2A*, differing only in the number of *CDKN2A* molecules detected per cell. *CDKN2A* status was called on the raw matrix counts, a cell counting as positive for an arm of the locus when it carried any non-zero count for that arm, and no further abundance threshold was applied. The distinct-UMI counts were used only for the recovery comparison against the matched whole-cell libraries. Each cell assayed by the capture was assigned a *CDKN2A* status from the *CDKN2A* transcripts detected in it, as p16^INK4a^-positive, p14^ARF^-positive, double-positive, positive for other *CDKN2A* transcripts only, or *CDKN2A*-negative. Cells whose barcode was not among the capture’s called cells were labelled not_in_pulldown and are distinguishable from measured zeros by a separate flag, and for the senescence contrasts they were treated as *CDKN2A*-negative. Capture cells were matched to the whole-cell atlas on donor and bare 10x barcode, both being in the same reverse-complement orientation.

Co-detection of p16^INK4a^ and p14^ARF^ was assessed only within cells called by the capture. Within each sample the number of double-positive cells expected under independence was computed as n × P(p16^INK4a^) × P(p14^ARF^), and observed and expected counts were summed within each sample and the log ratio was pooled across samples by random-effects meta-analysis with the Knapp–Hartung correction. Donors were treated as a second level by pooling samples within donor and summarising the four donor estimates with a one-sample t test. The null was computed with the depth stratifier collapsed to a single bin, so the reported enrichment is an upper bound. Cells not assayed by the capture carry no *CDKN2A* measurement and are excluded from this analysis entirely rather than treated as negatives.

### Definition of senescence-associated cells

Senescence-associated cells were defined as cells with a detected p16^INK4a^ transcript, comprising both p16^INK4a^-only, and p16^INK4a^/p14^ARF^ double-positive cells (n = 3,134). The comparator comprised all remaining cells not positive for *CDKN2A*, combining cells the capture assayed and in which no *CDKN2A* transcript was detected with cells the capture did not assay, which are treated as *CDKN2A*-negative as described above (n = 151,048). Cells positive only for p14^ARF^ (n = 23) or only for other *CDKN2A* transcripts (n = 912) were excluded from both arms, so that every senescence contrast is defined by p16^INK4a^ status alone. All senescence analyses are restricted to the four donors carrying a capture. Because a cell can be called p16^INK4a^-positive only if the capture recovered a *CDKN2A* molecule from it, and the capture selects for cells carrying more RNA, the senescence-associated arm is sequenced more deeply than the comparator. Depth is measured and matched on the annotated-ensemblid library size rather than on the gene-level total, except in the group-size analysis, where the match is on the ensemblid-level total.

### Depth matching for the senescence contrasts

Depth was controlled by matching on library size rather than by stratification, and the same procedure is shared by every senescence contrast reported except where stated otherwise. Cells were assigned to 20 equal-frequency bins of log_10_ library size. Within each bin every senescence-associated cell was retained and up to an equal number of comparator cells drawn at random without replacement; where a bin held fewer comparator cells than senescence-associated cells, all available comparators were taken, so the arms are not necessarily equal. Bins holding cells of only one arm contribute nothing. How much of the cohort survives depends on the contrast. The per-cell and per-gene contrasts retain all 3,134 senescence-associated cells against 2,963 comparator cells, 6,097 of 154,182 or 4.0% of the cohort; the 171-cell shortfall arises in depth bins holding fewer comparator cells than senescence-associated cells. The cell-class contrast, matched within class, retains 3,134 against 3,134, or 6,268 cells. The isoform-detection contrast is the one case in which senescence-associated cells are themselves discarded rather than merely left unpaired, retaining 2,931 against 2,931, because that contrast requires exactly balanced arms and a senescence-associated cell whose block holds no comparator cell at the same depth cannot be paired. Retention is quoted alongside the ratio. Matching is nested rather than pooled. For the per-cell, per-gene and isoform-detection contrasts it is performed within sample. Residual within-sample imbalance is measured rather than assumed and is reported per sample. For the cell-class-resolved contrast, matching is performed within broad cell class instead, for the reason given in that subsection. The per-cell and per-gene contrasts share a matched arm.

### Isoform detection at matched cell number and matched depth

Isoform detection was compared after equalizing both sequencing depth and cell number rather than on raw totals. Depth matching was applied first, as described above, which leaves 2,931 cells in each arm. The comparator arm was then subsampled to the senescence-associated cell count over 20 independent draws and the number of distinct isoforms detected were recorded per draw, overall and split by SQANTI3 structural category. The ratio reported is the mean over draws.

### Per-cell isoform diversity

Per-cell isoform diversity was calculated as the mean Shannon entropy of the isoform-fraction vectors of that cell’s genes, averaged over every gene the cell expresses, meaning every gene for which at least one isoform is detected in that cell. This unconditional denominator is the one used for every per-cell entropy result reported here. The senescent versus non-senescent contrast was estimated within sample on the depth-matched cells, with the sample (tube) as the unit of replication, requiring at least 5 cells per arm within a sample for that sample to contribute. No further depth stratification is applied within the matched arm. Per-sample effects were pooled by random-effects meta-analysis (DerSimonian–Laird) with the Knapp–Hartung small-sample correction, which is the primary result, and an assumption-free sign-flip permutation test is reported as the agreement check. Donor-level replication was assessed separately by pooling samples within donor and applying a one-sample t test to the four donor estimates. The same contrast was also run within each broad cell class on the matched cells, with a single stratum, on the classes carrying enough senescence-associated cells to estimate an effect.

### Per-gene changes in isoform diversity

For each gene, the difference in per-cell isoform entropy between senescence-associated and comparator cells was estimated within sample on the depth-matched cells, under the same unconditional denominator used for the per-cell metric, meaning that the mean is taken over all cells expressing the gene rather than only over those already detecting two or more of its isoforms. A gene was required to carry at least 5 cells per arm to be estimable, and p-values were Benjamini–Hochberg-corrected across genes.

The null was estimated empirically by permuting senescence status within sample 1,000 times and recomputing the whole statistic under each permutation, using the identical estimator applied to the observed data. The permutation null is computed on the depth-matched cells. The observed value is reported against this null as an excess and as a number of null standard deviations, and the raw percentage is not quoted on its own. Permutation p-values equal to 1/(n+1) are reported as the resolution floor rather than as measurements, and the z-score is quoted in preference where the floor is reached. A gene was called reproducible when it was estimable in at least 3 of the 4 donors and at least 3 of them agreed on the sign of the effect. Genes plotted in the volcano are the reproducible set, and the Benjamini–Hochberg correction is the full-set one, not re-run within the subset. The ratio between the arms of the fraction of expressing cells detecting two or more isoforms is reported as a check on the matching.

### Cell-class-resolved entropy changes and power

Per-gene entropy changes were re-estimated within each broad cell class carrying at least 50 senescence-associated cells, under the same unconditional denominator, requiring at least 5 cells per arm for a gene and class pair to be estimable. Classes below the threshold are reported as untested rather than dropped. Depth matching for this analysis was performed within each class rather than across the cohort. The within-class match uses the identical 20-bin equal-frequency procedure described above, applied before any accumulation, and per-class Cliff’s δ on library size is reported before and after matching.

Whether a non-significant gene and class pair reflects a genuine absence of effect or insufficient power was decided by computing, for each pair, the minimum detectable effect at 80% power and α = 0.05 from the observed within-class standard deviation and arm sizes. A pair was called genuinely null when its confidence interval excluded the effect measured in the class where the gene was significant, and underpowered otherwise. Robustness of each significant call to individual donors was assessed by leave-one-donor-out re-estimation, with three outcomes recorded, namely robust where all four re-estimates agree in sign, fragile where a sign reversal was observed, and unadjudicated where estimability was lost. Robustness percentages are summarized over the adjudicated calls, so that a call whose re-estimate was lost is not scored as a failure.

### Differential isoform usage in senescence

Differential isoform usage between senescence-associated and comparator cells was tested with the same gene-level Dirichlet-multinomial likelihood-ratio test described above, run independently within each broad cell class rather than pooled. Classes were required to carry a minimum number of senescence-associated cells pooled across donors and a minimum per-arm count within at least one donor. Apparent cell-type restriction was re-assessed on a common effect scale. For every gene called differentially used in at least one class, a single switch isoform was defined once, in that gene’s discovery class, meaning the class with the smallest adjusted p-value, as the isoform whose fraction changed most, and that same isoform was then read out in every class with at least 30 senescence-associated cells, requiring at least 5 cells per arm. A class was called genuinely null for that gene when its 95% confidence interval excluded the discovery-class effect, and uninformative or underpowered otherwise. Each gene’s effect elsewhere was additionally expressed as a fraction of its own discovery-class effect. In parallel, every class was downsampled to equal senescence-associated cell counts of 50 and 100, with four reference cells per senescence-associated cell and three replicates each, and the restriction distribution recomputed.

Unlike the diversity contrasts above, this analysis is not depth-controlled. The depth stratifier is collapsed to a single bin, so the contrast runs on all cells of a class rather than on a depth-matched subsample, and the estimates are reported as an upper bound on how much cell-class structure survives.

### Statistics and reporting conventions

Analyses were performed in Python (3.11.13 and 3.12.12) using scanpy v1.11.5, anndata v0.12.7, numpy v2.3.5, scipy v1.17.0, pandas v2.3.3, scikit-learn v1.8.0 and statsmodels v0.14.6. Pipelines were orchestrated with Snakemake v7.32.4 on a Slurm cluster. Unless stated otherwise, multiple testing was controlled by the Benjamini–Hochberg procedure and significance thresholds are stated at each point of use. Correlations are Spearman unless Pearson is named.

For the senescence analyses, where cells within a sample and within a donor are not independent, a fixed set of conventions was applied throughout. The sample (tube) is the primary unit of replication, pooled by random-effects meta-analysis with the Knapp–Hartung correction. Donor-level replication is summarized by a one-sample t test on the four donor estimates. Effect sizes with confidence intervals are reported in preference to p-values.

Sequencing depth differs systematically between the senescence-associated and comparator arms, so every contrast states its depth control explicitly. The isoform-detection, per-cell entropy, per-gene entropy and cell-class-resolved contrasts are depth-matched as described above, and the per-cell and per-gene entropy metrics use the unconditional denominator, meaning that the mean is taken over every cell expressing a gene rather than only over cells already detecting two or more of its isoforms. The *CDKN2A* co-detection statistic and the differential isoform usage analysis are not depth-matched, and both are therefore upper bounds on the effect attributable to senescence status. Where an estimate is quoted without a stated depth control, it is depth-matched.

