## Supplementary File 1 for "Single-cell splice isoform usage reveals distinct axes of cellular identity and senescence"

### Supplementary Figures

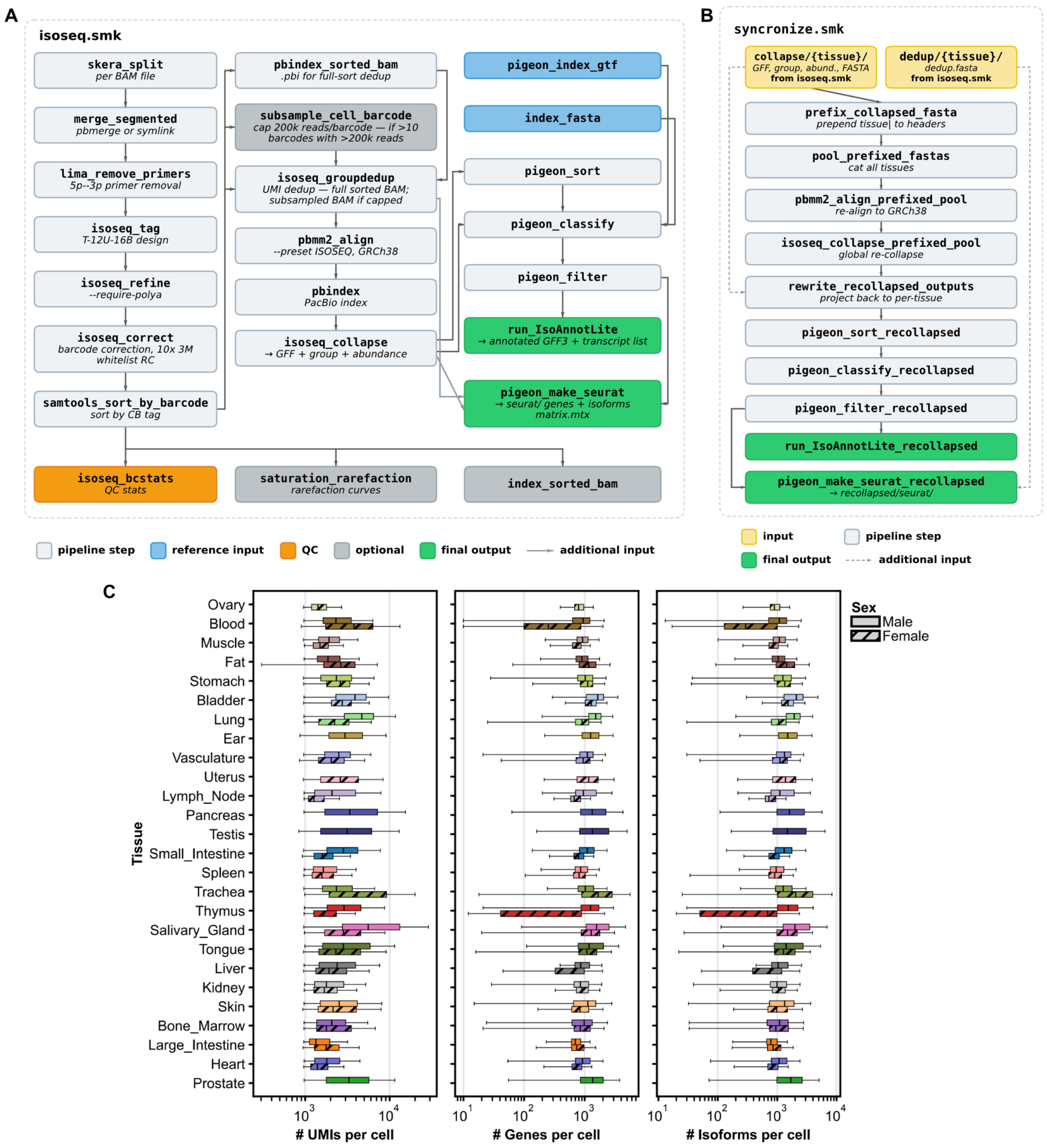


**Supplementary Figure 1. Processing pipeline and cross-tissue harmonization.**
**(A)** Snakemake rule graph for the Iso-Seq pipeline, the per-tissue Iso-Seq module. Long-read BAMs are deconcatenated (skera_split), merged, and stripped of 5'/3' primers (lima_remove_primers). Cell barcodes and UMIs are extracted under a T-12U-16B read design (isoseq_tag), polyA-containing reads are retained (isoseq_refine), and barcodes are corrected against the 10x whitelist (isoseq_correct). Barcode-sorted reads are UMI-deduplicated (isoseq_groupdedup), aligned to GRCh38 (pbmm2_align), and collapsed into per-tissue transcript models (isoseq_collapse). Models are classified and filtered against the reference annotation with Pigeon, yielding an IsoAnnotLite-annotated GFF3 and a Seurat-compatible isoform × cell matrix. Node color indicates rule type; dashed edges indicate additional inputs to a rule.
**(B)** Snakemake rule graph for the PacBio identifier synchronization pipeline. Per-tissue collapsed FASTAs are prefixed with their tissue of origin, pooled, re-aligned to GRCh38, and globally re-collapsed so that transcript identifiers are consistent across tissues. Recollapsed models are projected back to per-tissue coordinates and passed through the same Pigeon sort–classify–filter sequence as in A.
**(C)** Per-cell library statistics by tissue and sex: UMIs per cell (left), genes detected per cell (middle), and isoforms detected per cell (right), all on a $\log_{10}$ scale. Boxes span the interquartile range with the median marked; whiskers extend to 1.5 × IQR. Color encodes tissue and hatching encodes donor sex (solid, male; hatched, female). Tissues are ordered by total cell number as in Figure 1B.

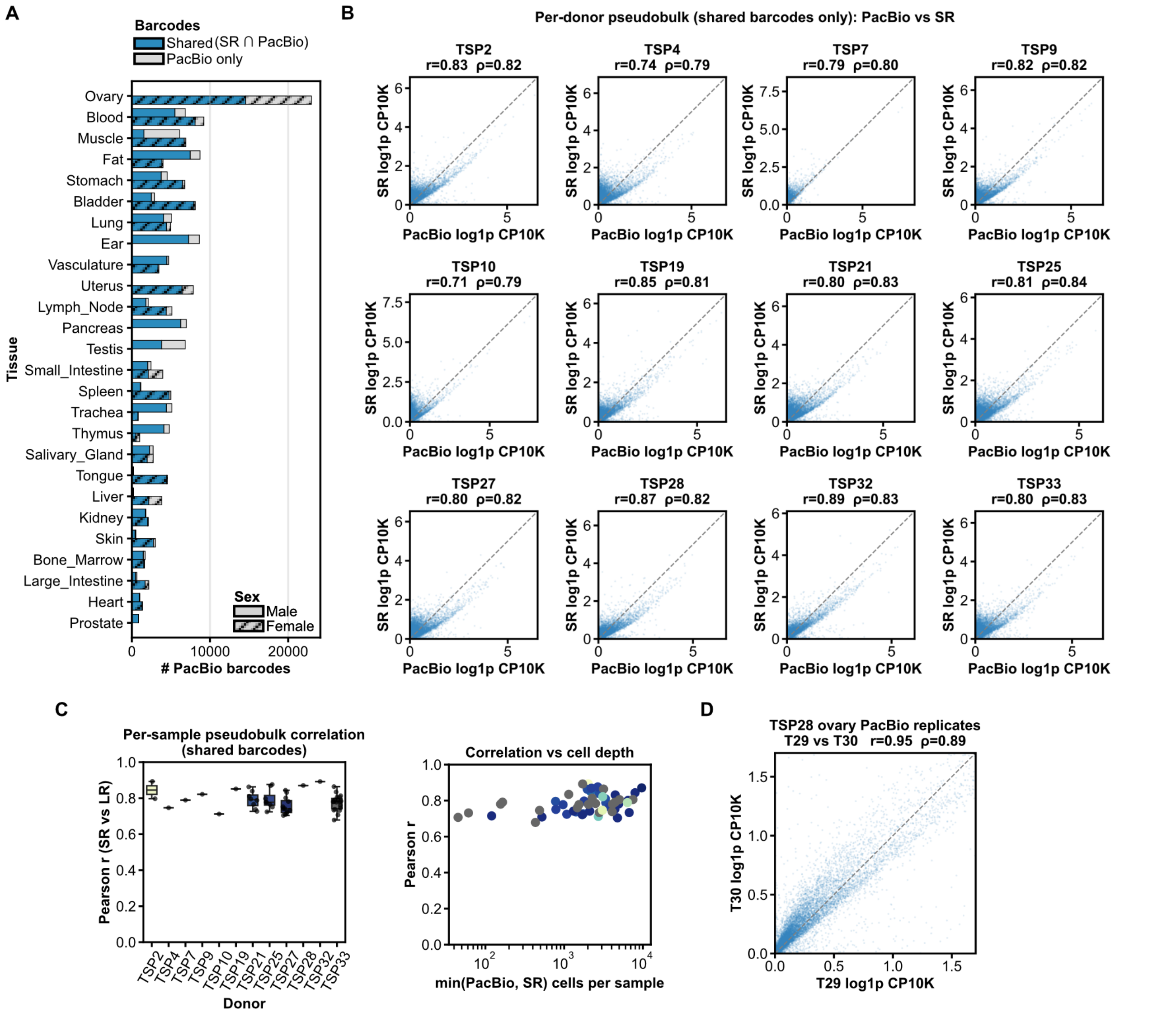


**Supplementary Figure 2. Concordance between matched long-read and short-read measurements.**
**(A)** Concordance between short-read (SR) and long-read cell barcodes per tissue. Bar plot showing the number of long-read cell barcodes per tissue, partitioned into those also recovered in the matched short-read library (shared, blue) and those unique to long-read (grey); hatching encodes donor sex as in Supp. Figure 1C.
**(B)** Per-donor pseudobulk comparison of long-read and short-read (SR) expression, restricted to cell barcodes recovered in both assays. For each donor, counts were summed across shared barcodes, normalized to counts per 10,000 (CP10K), and log1p-transformed; each point is one gene (24,979–41,251 genes per donor). The dashed line is the identity. Pearson (r) and Spearman (ρ) correlations are given above each panel.
**(C)** Distribution of per-sample long-read versus short-read gene expression correlations across matched samples.
**(D)** Long-read technical reproducibility: pseudobulk gene expression for two replicate libraries prepared from the same tissue, T29 versus T30.

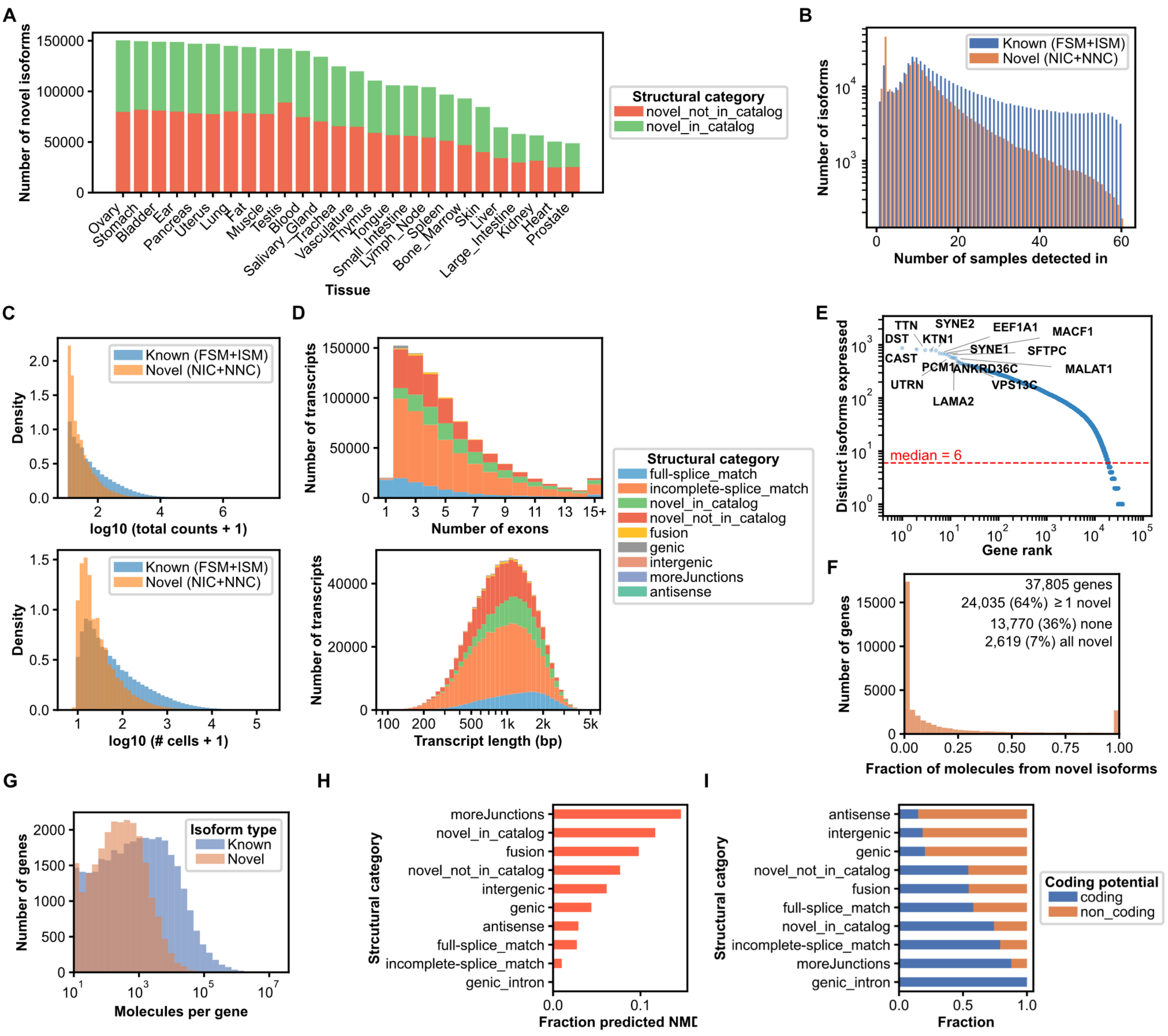


**Supplementary Figure 3. Properties of novel isoforms across tissues.**

**(A)** Number of novel isoforms per tissue, split into novel-in-catalog (NIC) and novel-not-in-catalog (NNC). Tissues are ordered by novel-isoform count.

**(B)** Number of samples in which each novel isoform is detected (>= 1 count in >= 1 cell of that sample), across all 60 samples. The y-axis is log-scaled.

**(C)** Expression and detection properties of known (FSM+ISM) versus novel (NIC+NNC) isoforms. Top: density of isoforms by expression level, $\log_{10}$(total counts + 1). Bottom: density by detection breadth, $\log_{10}$(number of cells + 1).

**(D)** Structural composition as a function of transcript complexity, colored by structural category (legend at right). Top: number of transcripts by exon count (1 to 15+). Bottom: number of transcripts by transcript length (bp).

**(E)** Number of distinct isoforms expressed per gene, with genes ordered by rank. Both axes are log scaled. The dashed red line marks the median of 6 isoforms per gene. Selected genes at the high-diversity end of the distribution are labelled.

**(F)** Distribution across genes of the fraction of each gene's molecules contributed by novel (NIC + NNC) isoforms; y = number of genes. Every gene with at least one molecule is included, so the leftmost bar is the genes with no novel molecules and the rightmost the genes whose molecules are entirely novel. Dashed line, mean across genes; dotted line, the pooled molecule-weighted fraction across all genes.

**(G)** Distribution across genes of molecule counts, split by isoform type: known (FSM + ISM) versus novel (NIC + NNC); log-spaced bins, y = number of genes. Genes contributing no molecules of a class are omitted from that histogram.

**(H)** Fraction of isoforms predicted to be nonsense-mediated decay (NMD) targets within each structural category, ordered by increasing NMD rate (bottom to top).

**(I)** Fraction of isoforms predicted to be coding versus non-coding within each structural category, ordered by increasing coding fraction (top to bottom).

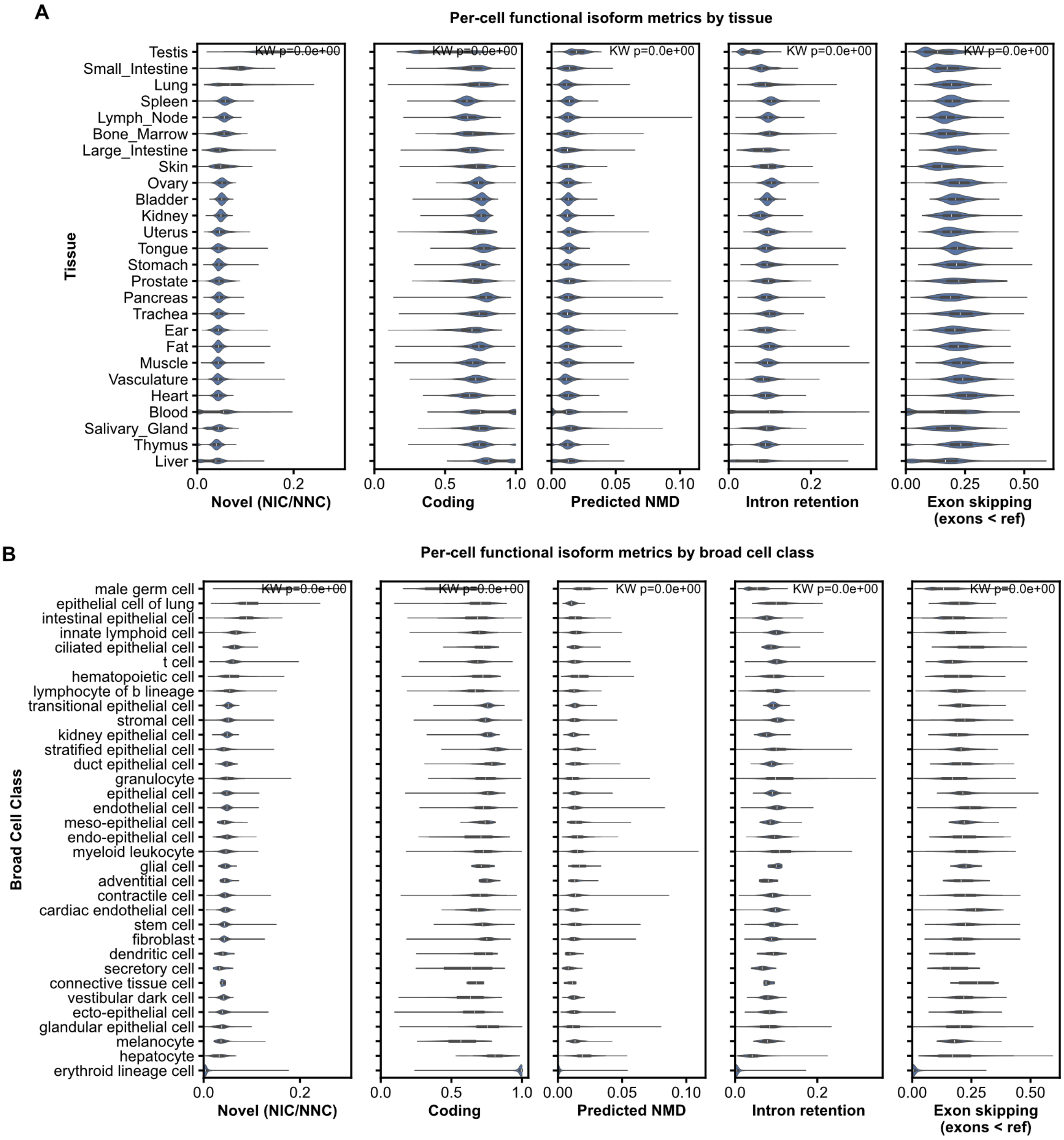


**Supplementary Figure 4. Functional isoform properties across tissues and cell types.**
**(A)** Per-cell distributions of novel-splicing, coding-potential, nonsense-mediated decay, intron-retention and exon-skipping fractions across tissues.
**(B)** The same per-cell metrics across broad cell classes.

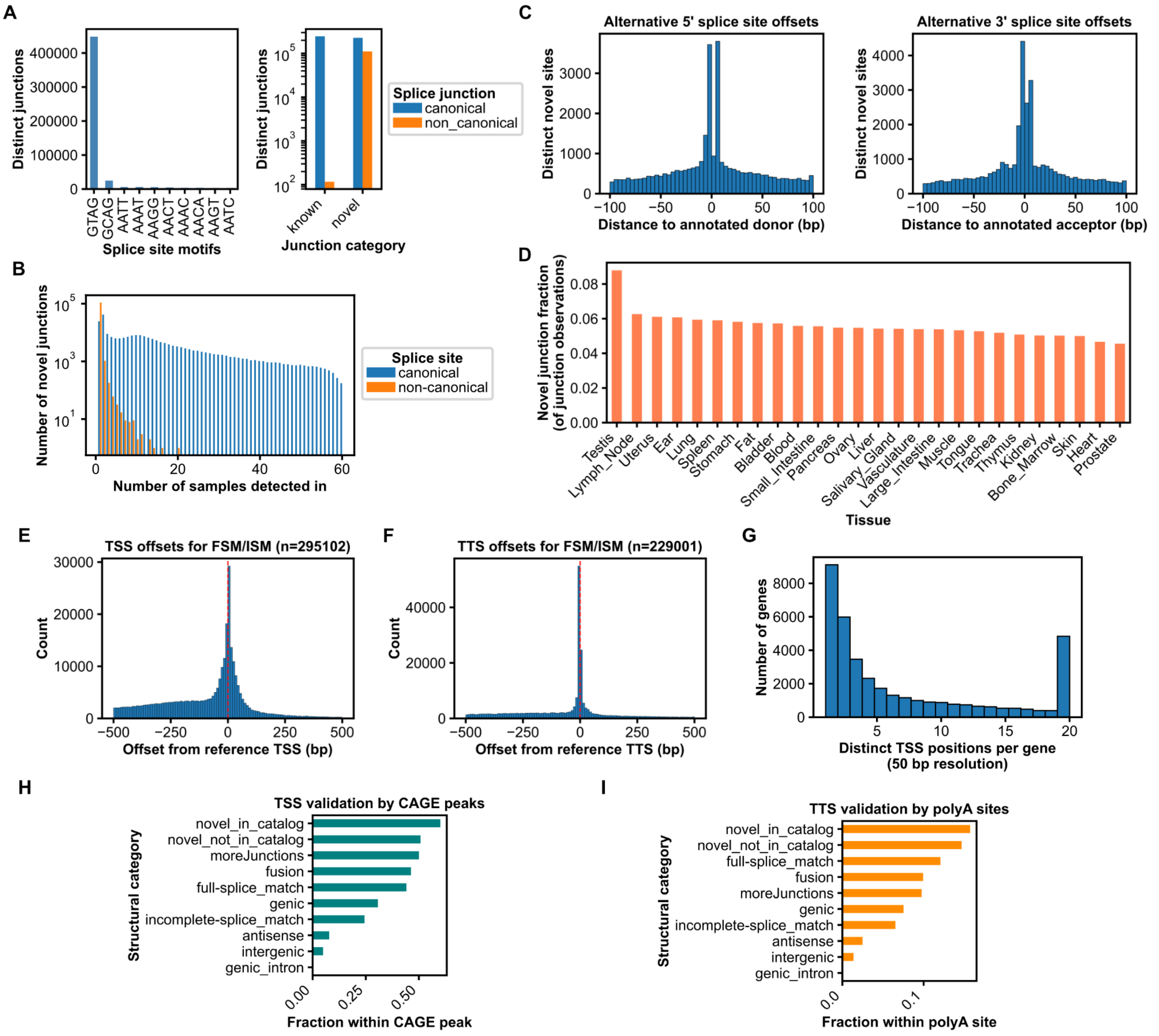


**Supplementary Figure 5. Splice-junction and transcript-boundary quality control for long-read transcript models relative to a reference annotation.**

**(A)** Splice donor–acceptor motif usage. Left, counts of dinucleotide motifs across all splice junctions; the canonical GT–AG motif vastly outnumbers all others combined. Right, canonical (blue) vs. non-canonical (orange) junction counts split by annotation status (known vs. novel; log-scaled y-axis).

**(B)** Number of novel splice junctions, by canonical (blue) and non-canonical (orange) status, as a function of the number of samples each junction was independently detected in (log-scaled y-axis).

**(C)** Distance (bp) between each alternative 5' (left) and 3' (right) splice site and its nearest annotated counterpart.

**(D)** Fraction of splice junctions per tissue that are novel relative to the reference.

**(E)** Offset (bp) between observed and reference transcription start sites (TSS; n = 295,102), restricted to full-splice-match and incomplete-splice-match transcripts; dashed red line marks zero offset.

**(F)** Offset (bp) between observed and reference transcription termination sites (TTS; n = 229,001), plotted as in (E).

**(G)** Number of distinct TSS positions per gene at 50 bp resolution. Counts are capped at 20; the rightmost bar pools all genes with 20 or more distinct TSS positions.

**(H)** Independent validation of transcript 5' ends by structural category: fraction of transcripts with a TSS inside a CAGE peak.

**(I)** Independent validation of transcript 3' ends by structural category: fraction of transcripts with a TTS inside an annotated polyA site, plotted as in (H).

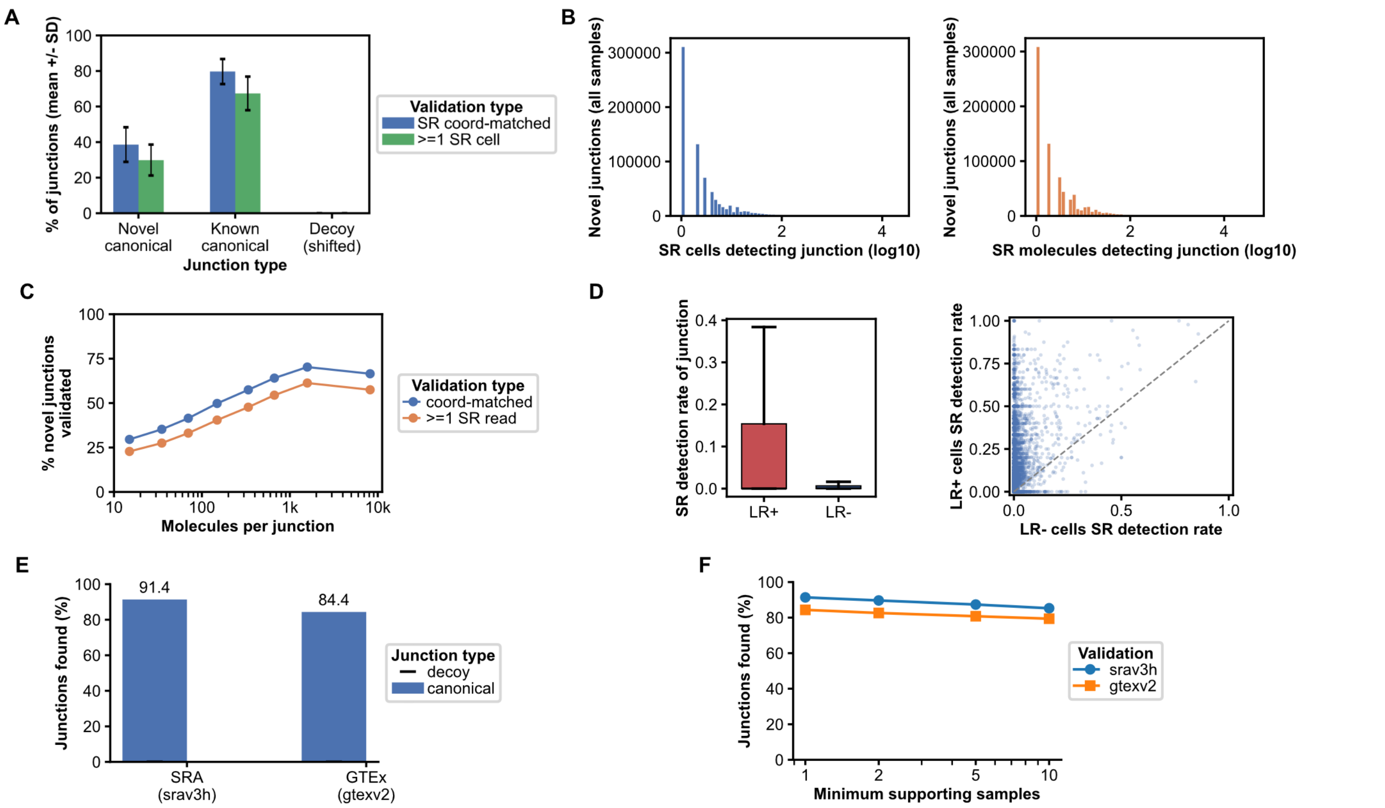


**Supplementary Figure 6. Short-read validation of novel splice junctions.**

**(A)** Per-sample validation rate for novel junctions, known junctions, and coordinate-shifted decoy junctions, across 48 matched short-read samples.

**(B)** Short-read support depth for validated junctions, shown as supporting cells and supporting molecules.

**(C)** Validation rate as a function of long-read expression, binned by the number of molecules supporting each junction.

**(D)** Per-cell validation: short-read detection of a novel junction in cells that express its isoform in long-read data (LR+) versus cells that express the gene without that isoform (LR-).

**(E)** Fraction of novel junctions recovered in the Snaptron SRA (srav3h) and GTEx (gtexv2) compilations. Black dashes mark the coordinate-shifted decoy rate for the same junction set.

**(F)** Fraction of novel canonical junctions recovered as a function of the minimum number of Snaptron samples required to count a junction as found, for each compilation.

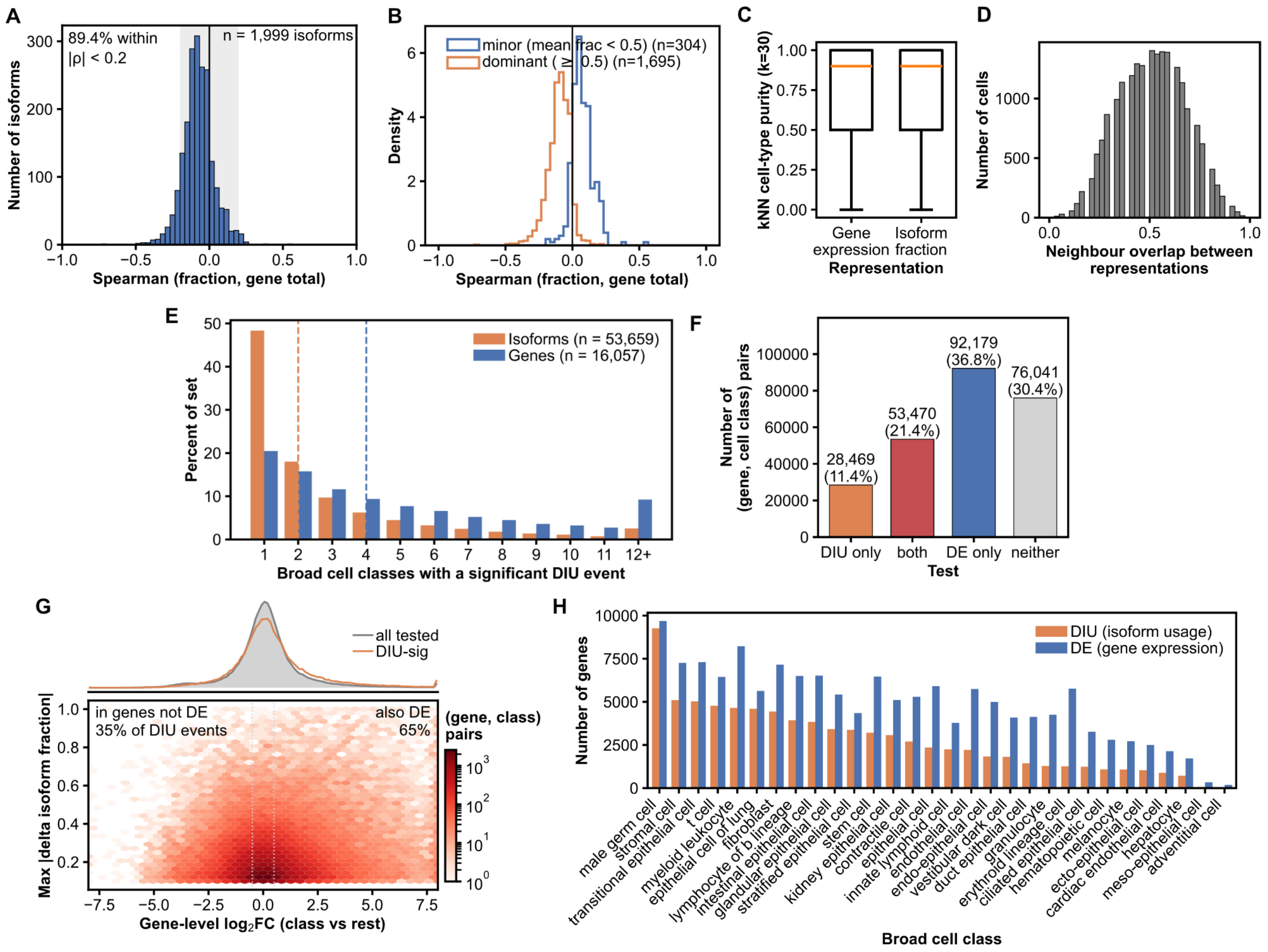


**Supplementary Figure 7. Isoform usage as an axis of cell identity distinct from gene expression.**
**(A)** Distribution of the Spearman correlation between an isoform's fraction and its gene's total count, across the isoform features used for the embedding. The shaded band marks |rho| < 0.2, which contains 89.4% of isoforms.
**(B)** The Spearman correlation between an isoform's fraction and its gene's total count, as in (A), split by isoform dominance. Isoforms are split at a mean fraction of 0.5 into minor and dominant classes. The effect is small in both directions.
**(C)** Distribution of per-cell k-nearest-neighbour cell-type purity (k = 30) for cells embedded on gene expression versus on isoform fractions, with both representations restricted to the same set of genes. Boxes span the interquartile range with the median marked.
**(D)** Overlap between each cell's k nearest neighbours in the gene-expression space and in the isoform-fraction space. The dashed line marks the median (0.50).
**(E)** Number of broad cell classes in which each significant isoform, and each significant gene, shows differential isoform usage, as a percent of each set. Dashed lines mark the two medians; the last bar aggregates 12 or more classes.
**(F)** Four-way partition of gene and cell class pairs by whether the signal lies in isoform usage only, in gene expression only, in both, or in neither. Bars are annotated with counts and percentages.
**(G)** Joint density of gene-level $\log_2$ fold change against the largest isoform fraction shift, restricted to pairs with significant differential isoform usage. The grey marginal shows the fold-change distribution of all tested pairs for reference.
**(H)** Number of genes with differential isoform usage and number of differentially expressed genes, per broad cell class. Both counts are taken from the same set of testable pairs, so the two bars share a single denominator.

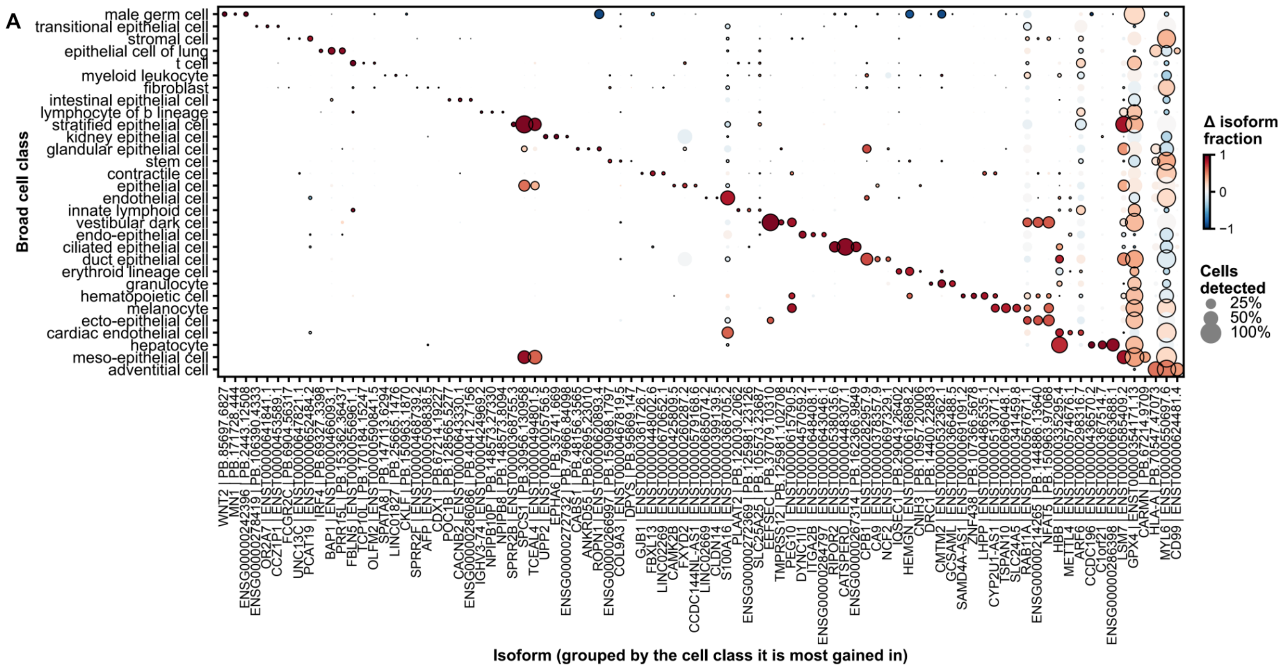


**Supplementary Figure 8. Differential isoform usage across broad cell classes.**
**(A)** The three isoforms most strongly gained in each broad cell class (isoforms with a positive change in isoform fraction only; each isoform assigned to the single class in which its gain is largest). Dot color shows the change in isoform fraction between that class and all other cells, and dot size shows the fraction of cells expressing the gene.

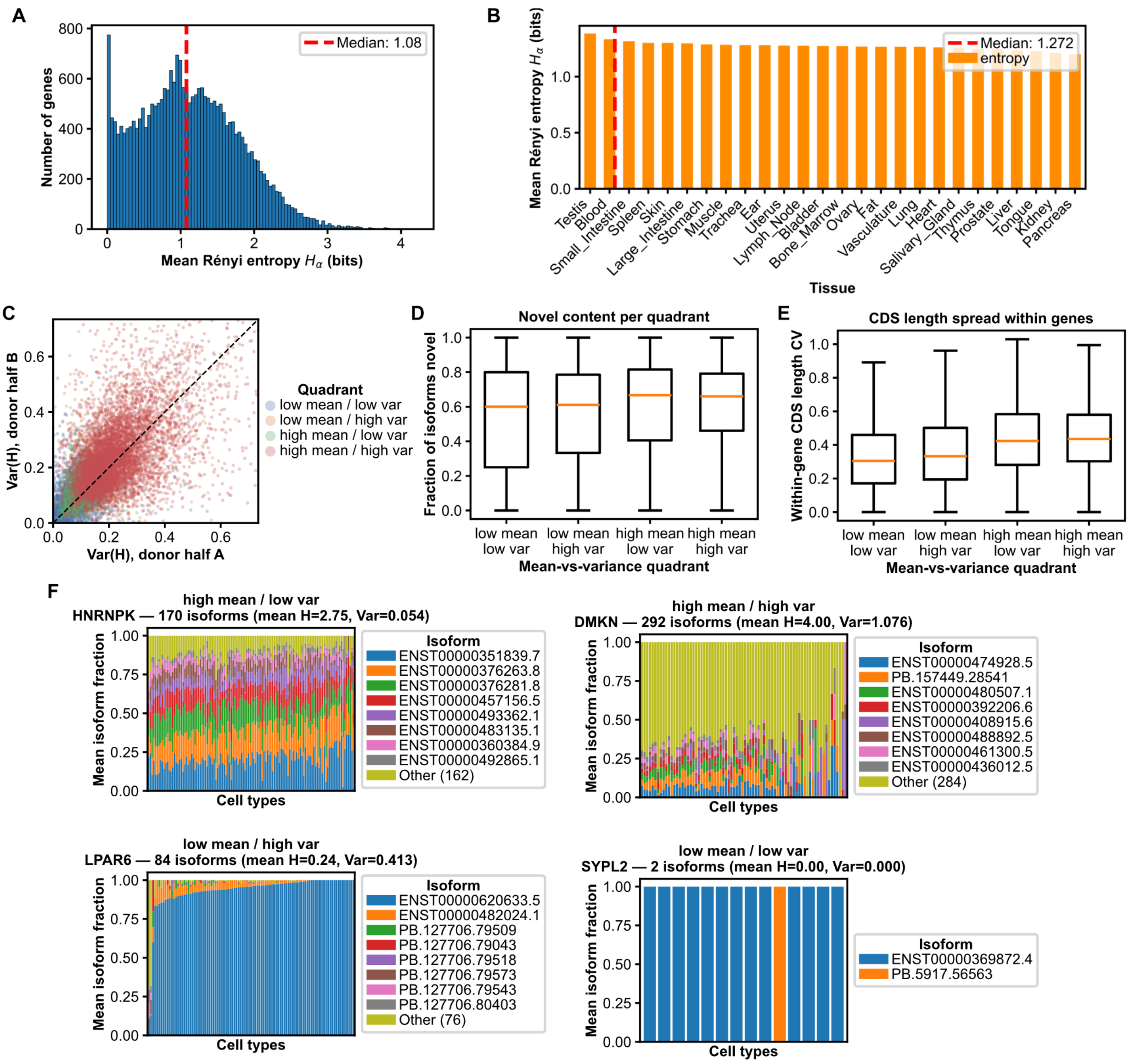


**Supplementary Figure 9. Distribution of isoform diversity across genes, cell types and tissues.**
**(A)** Distribution of per-gene Rényi entropy, each gene summarized as its mean across the cell types in which it was tested (26,885 genes). The dashed line marks the median.
**(B)** Mean per-gene Rényi entropy per tissue, ordered by decreasing entropy.
**(C)** Split-half reproducibility between donors of the across-cell-type variance of Rényi entropy, the quantity that defines the quadrants in Figure 3F.
**(D)** Fraction of each gene's isoforms that are novel (NIC or NNC), by mean-versus-variance quadrant.
**(E)** Coefficient of variation of coding-sequence length among the coding isoforms of the same gene, by mean-versus-variance quadrant. Genes with at least two coding isoforms.
**(F)** Isoform usage of representative genes drawn from each mean-versus-variance quadrant, shown as stacked isoform fractions across cell types.

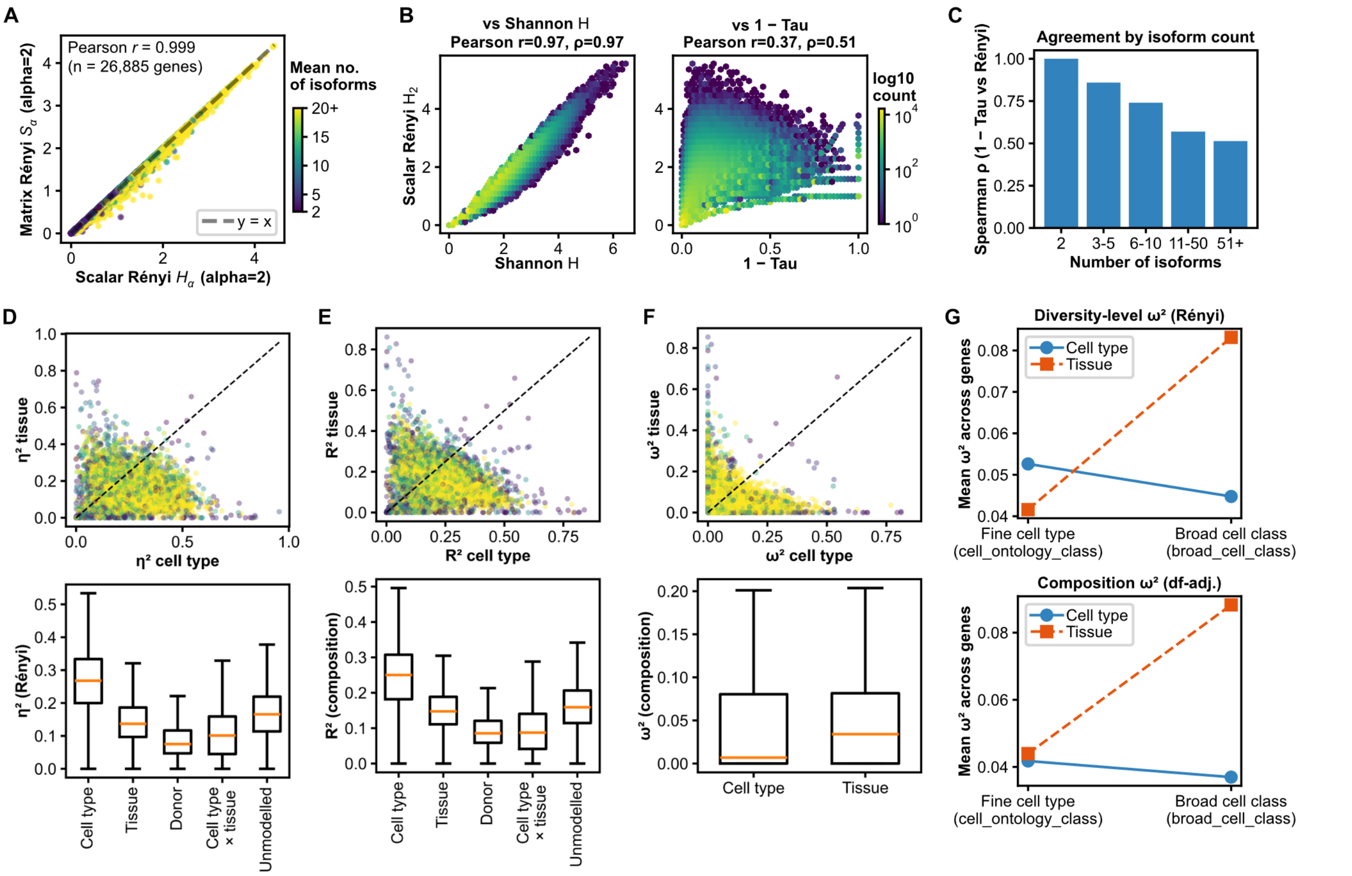


**Supplementary Figure 10. Robustness of the diversity metrics to the cell-type-versus-tissue partition.**
**(A)** Scalar Rényi entropy versus the matrix-based Rényi entropy that incorporates per-cell heterogeneity. Both entropies are averaged within a gene across the cell types in which it is expressed, so each point is one gene (n = 26,885); points are colored by the gene's mean number of detected isoforms, and the dashed line is the identity.
**(B)** Scalar Rényi entropy versus Shannon entropy and versus 1 – Tau, at the gene-by-cell-type level.
**(C)** Relationship between 1 – Tau and Rényi entropy, stratified by the number of isoforms per gene.
**(D)** Raw $\eta^2$ for isoform diversity attributable to cell type versus tissue, one point per gene (left), and the distribution across genes of all five terms, namely cell type, tissue, donor, the cell type × tissue interaction and unmodelled variation (right). "Unmodelled" is what remains after the interaction is removed and is not the additive model's residual. Raw $\eta^2$ is not degrees-of-freedom adjusted and mechanically favours the factor with more levels.
**(E)** Raw PERMANOVA $R^2$ for isoform composition attributable to cell type versus tissue, one point per gene (left), and the distribution of $R^2$ across genes for all five terms, namely cell type, tissue, donor, the cell type × tissue interaction and unmodelled variation (right), defined as in (D). Raw $R^2$ is not degrees-of-freedom adjusted and mechanically favours the factor with more levels.
**(F)** Degrees-of-freedom-adjusted $\omega^2$ for isoform composition attributable to cell type and to tissue: per-gene scatter (left) and distribution across genes (right).
**(G)** Resolution ladder: mean $\omega^2$ attributable to cell type and to tissue as the cell-type annotation is coarsened from 144 fine cell types to 34 broad classes, for both the composition and diversity-level arms.

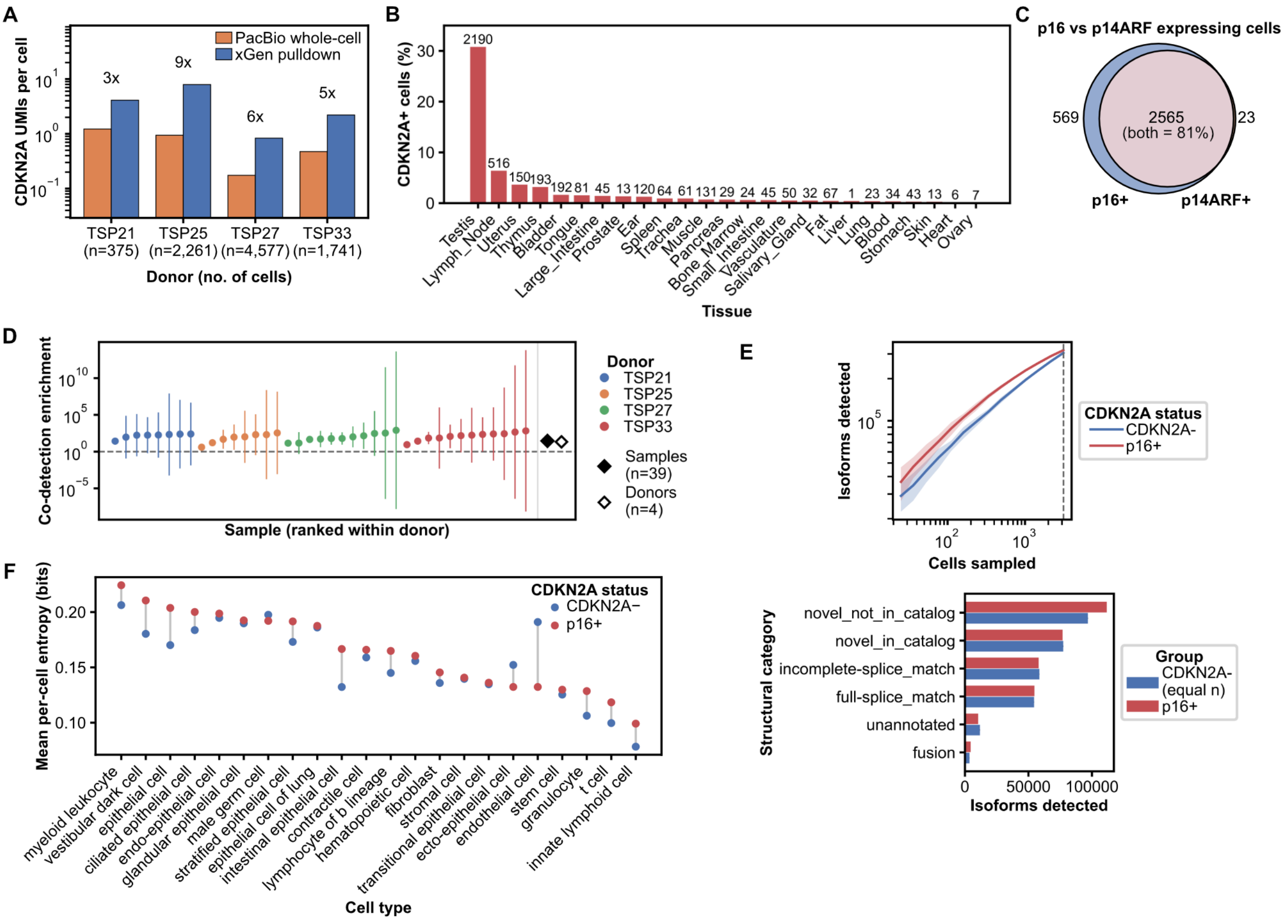


**Supplementary Figure 11. Senescent cell identification and isoform diversity across cell types.**
**(A)** Mean distinct *CDKN2A* UMIs recovered per cell by the targeted capture versus the matched long-read whole-cell library, per donor, on a log scale with fold change annotated. Restricted to the 8,954 cells the capture assayed.
**(B)** Percentage of cells called *CDKN2A*-positive by the capture, per tissue.
**(C)** Overlap of cells detecting p16$^{INK4a}$ and p14$^{ARF}$ isoforms in the capture: 569 p16$^{INK4a}$-positive only, 23 p14$^{ARF}$-positive only, and 2,565 detecting both.
**(D)** Per-sample enrichment of p16$^{INK4a}$/p14$^{ARF}$ co-detection over an empirical null, with pooled sample- and donor-level estimates.
**(E)** Top, rarefaction of isoform discovery, with distinct isoforms detected as a function of the number of cells sampled, separately for p16$^{INK4a}$-positive and *CDKN2A*-negative cells. Bottom, isoforms detected per SQANTI3 structural category in the same two groups at matched cell number. Cells are 1:1 matched on library size between the groups throughout, so both panels differ in isoform discovery and not in sequencing depth.
**(F)** Mean per-cell isoform entropy within each broad cell class, on library-size-matched cells. Each pair of points is one class, joined by a connector whose length is that class's effect; blue, *CDKN2A*-negative; red, p16$^{INK4a}$-positive. Classes are ordered by their p16$^{INK4a}$-positive mean.

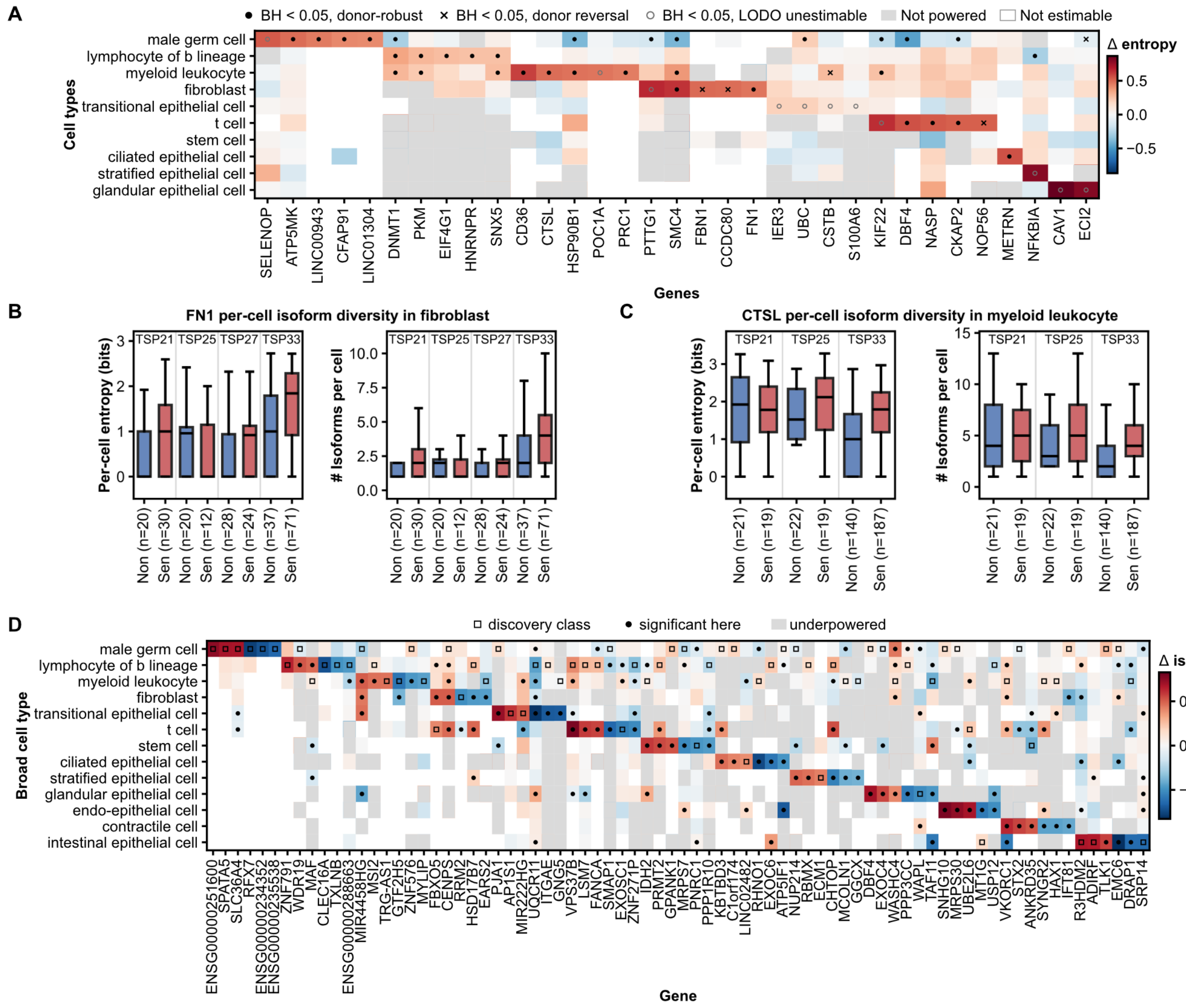


**Supplementary Figure 12. Cell-type-resolved isoform entropy changes in p16$^{INK4a}$-positive cells, and the power to detect them.**

**(A)** Change in per-cell isoform entropy for up to five genes per broad cell class, selected among genes whose entropy increases in that class, shown in every class. Filled dot, BH < 0.05 and robust to leaving out any donor; cross, BH < 0.05 but a sign reversal was observed when a donor was left out (5); open circle, BH < 0.05 but leave-one-donor-out could not adjudicate the call because estimability was lost (11). Grey, class not powered to detect the effect seen where it was significant. White, not estimable.

**(B)** Per-cell isoform diversity for *FN1* in fibroblasts: entropy in bits (left) and isoforms detected per cell (right), by donor and *CDKN2A* status. Entropy is averaged over every cell expressing the gene rather than only those already using multiple isoforms, so the two panels describe the same set of cells.

**(C)** The same view for *CTSL* in myeloid leukocytes, a representative comparison highlighted in panel A: per-cell isoform entropy (left) and isoforms detected per cell (right), by donor and *CDKN2A* status.

**(D)** Change in isoform fraction for the top five differential isoform usage switches of each broad cell class, shown across all classes. Columns are the union of the selected gene and isoform pairs and rows are broad cell classes, with color indicating the change in isoform fraction between p16$^{INK4a}$-positive and *CDKN2A*-negative cells. Open square, the class in which the switch was defined, which is not independent evidence of an effect there; filled dot, independently significant in that class; grey, class not powered to detect the effect seen in the discovery class; white, not estimable.
