## Supplementary File 2 for "Single-cell splice isoform usage reveals distinct axes of cellular identity and senescence"

**Full author list and affiliations in Tabula Sapiens Consortium:**


Robert C. Jones[1]
Mark A. Krasnow[2,3]
Angela Oliveira Pisco[4]
Stephen R. Quake[1,5,6]
Julia Salzman[2,7]
Nir Yosef[4,8,9]
Madhav Mantri[1]
Anton Thieme[1]
Jessie Aguirre[10]
Ron Garner[10]
Sal Guerrero[10]
William Harper[10]
Resham Irfan[10]
Sophia Mahfouz[10]
Ravikumar Ponnusamy[10]
Bhavani A. Sanagavarapu[10]
Ahmad Salehi[10]
Ivan Sampson[10]
Chloe Tang[10]
Alan G. Cheng[11]
James M. Gardner[12,13]
Burnett Kelly[10,14]
Lindsay Moore[15]
Thurman Slone[10]
Zifa Wang[10]
Anika Nawar Choudhury[1,4]
Sheela Crasta[1]
George Crowley[1]
Chen Dong[1,4]
Marcus L. Forst[1]
Doug E. Henze[1]
Jaeyoon Lee[1]
Maurizio Morri[4]
Serena Y. Tan[16]
Sevahn K. Vorperian[17,18]
Lynn T. Yang[1,4]
Marcela Alcántara-Hernádez[19]
Julian Berg[20]
Dhruv Bhatt[21]
Sara Billings[11]
Andrès Gottfried-Blackmore[19,22]
Jamie Bozeman[21]
Simon Bucher[23]
Elisa Caffrey[24]
Amber Casillas[25]
Rebecca Chen[24]
Matthew Choi[23]
Rebecca N. Culver[26]
Ivana Cvijovic[1,5]

Ke Ding[27]
Hala Shakib Dhowre[28]
Hua Dong[27]
Kenneth Donaville[23]
Lauren Duan[20]
Xiaochen Fan[21]
Mariko H. Foecke[29]
Francisco X. Galdos[20]
Eliza A. Gaylord[29]
Karen Gonzales[21]
William R. Goodyer[30]
Michelle Griffin[31]
Yuchao Gu[2,32,33]
Shuo Han[27]
Jun Yan He[24]
Paul Heinrich[20]
Rebeca Arroyo Hornero[19]
Keliana Hui[24]
Juan C. Irwin[25]
SoRi Jang[2]
Annie Jensen[20,34]
Saswati Karmakar[16,26]
Jengmin Kang[35]
Hailey Kang[23]
Bikem Soygur[29]
Soochi Kim[35]
Stewart J. Kim[2,32,33]
William Kong[27]
Mallory A. Laboulaye[27]
Daniel Lee[20]
Gyehyun Lee[36]
Elise Lelou[23]
Anping Li[27]
Baoxiang Li[28]
Wan-Jin Lu[27]
Hayley Raquer-McKay[19]
Elvira Mennillo[36]
Elena Montauti[24]
Karim Mrouj[27]
Shravani Mukherjee[28]
Patrick Neuhöfer[2,32,33]
Saphia Nguyen[23]
Honor Paine[23]
Jennifer B. Parker[27,31]
Julia Pham[24]
Kiet T. Phong[37]
Pratima Prabala[21]
Zhen Qi[27]
Joshua Quintanilla[20,34]
Iulia Rusu[36]
Alireza Raissadati[20]

Bronwyn Scott[28]
David Seong[19]
Hosu Sin[38]
Hanbing Song[39]
Sean P. Spencer[19,22]
Varun R. Subramaniam[28]
Michael Swift[1]
Aditi Swarup[28]
Greg Szot[12,13]
Aris Taychameekiatchai[23]
Emily Trimm[21]
Stefan Veizades[20,34]
Sivakamasundari Vijayakumar[27]
Kim Chi Vo[25]
Tian Wang[40]
Timothy T. H. Wu[2]
Yinghua Xie[20,34]
William Yue[23]
Yue Zhang[2]
Angela Detweiler[4]
Honey Mekonen[4]
Norma F. Neff[4]
Sheryl Paul[4]
Amanda Seng[4]
Jia Yan[4]
Deana Rae Crystal Colburg[41]
Balint Laszlo Forgo[16]
Luca Ghita[19]
Frank McCarthy[42]
Aditi Agrawal[4]
Alina Isakova[1]
Kavita Murthy[1]
Alexandra Psaltis[1]
Wenfei Sun[1]
Kyle Awayan[4]
Pierre Boyeau[43]
Robrecht Cannoodt[44–46]
Leah Dorman[4]
Samuel D'Souza[4]
Can Ergen[8,43]
Siyu He[1,7]
Justin Hong[47]
Harper Hua[7]
Erin McGeever[4]
Antoine de Morree[35,48]
Luise A. Seeker[1]
Alexander J. Tarashansky[4]
Astrid Gillich[2]
Taha A. Jan[49]
Angela Ling[49]
Abhishek Murti[23]

Nikita Sajai[23]
Ryan M. Samuel[50]
Juliane Winkler[51,52]
Steven E. Artandi[2,32,33]
Philip A. Beachy[27,34,53]
Mike F. Clarke[27]
Zev Gartner[4,54]
Linda C. Giudice[25]
Franklin W. Huang[39,55]
Juliana Idoyaga[19,56]
Michael G. Kattah[36]
Christin S. Kuo[57]
Diana J. Laird[29]
Michael T. Longaker[27,58]
Patricia Nguyen[20,34,59]
David Y. Oh[24]
Thomas A. Rando[35]
Kristy Red-Horse[21]
Bruce Wang[23]
Albert Y. Wu[28]
Sean M. Wu[20,34]
Bo Yu[38]
James Zou[7,60]

**Affiliations**

1. Department of Bioengineering, Stanford University; Stanford, CA, USA.
2. Department of Biochemistry, Stanford University School of Medicine, Stanford, CA, USA.
3. Howard Hughes Medical Institute, USA.
4. Chan Zuckerberg Biohub, San Francisco, CA, USA.
5. Department of Applied Physics, Stanford University, Stanford, CA, USA.
6. The Chan Zuckerberg Initiative, Redwood City, CA, USA.
7. Department of Biomedical Data Science, Stanford University, Stanford, CA, USA.
8. Center for Computational Biology, University of California Berkeley, Berkeley, CA, USA.
9. Ragon Institute of MGH, MIT and Harvard, Cambridge, MA, USA.
10. Donor Network West, San Ramon, CA, USA.
11. Department of Otolaryngology-Head and Neck Surgery, Stanford University School of Medicine, Stanford, California, USA.
12. Department of Surgery, University of California San Francisco, San Francisco, CA, USA.
13. Diabetes Center, University of California San Francisco, San Francisco, CA, USA.
14. DCI Donor Services, Sacramento, CA, USA.
15. Division of Neurotology and Lateral Skull Base Surgery Department of Otolaryngology- Head & Neck Surgery Stanford University, Stanford, CA, USA.
16. Department of Pathology, Stanford University School of Medicine, Stanford, CA, USA.
17. Department of Chemical Engineering, Stanford University, Stanford, CA, USA.
18. Sarafan ChEM-H, Stanford University, Stanford, CA, USA.
19. Department of Microbiology and Immunology, Stanford University School of Medicine, Stanford, CA, USA.
20. Stanford Cardiovascular Institute, Stanford CA, USA.
21. Department of Biology, Stanford University, Stanford, CA, USA.
22. Division of Gastroenterology, Department of Medicine, Stanford University School of Medicine, Stanford, CA, USA.

23. Department of Medicine and Liver Center, University of California San Francisco, San Francisco, CA, USA.
24. Division of Hematology/Oncology, Department of Medicine, University of California San Francisco, San Francisco, CA, USA.
25. Center for Reproductive Sciences, Department of Obstetrics, Gynecology and Reproductive Sciences, University of California San Francisco, San Francisco, CA, USA.
26. Department of Genetics, Stanford University School of Medicine, Stanford, CA, USA.
27. Institute for Stem Cell Biology and Regenerative Medicine, Stanford University School of Medicine, Stanford, CA, USA.
28. Department of Ophthalmology, Stanford University School of Medicine, Stanford, CA, USA.
29. Department of Ob/Gyn and Reproductive Sciences, Eli and Edythe Broad Center for Regeneration Medicine and Stem Cell Research, University of California, San Francisco.
30. Department of Pediatrics, Division of Cardiology, Stanford University School of Medicine, Stanford, CA, USA.
31. Department of Surgery, Division of Plastic and Reconstructive Surgery, Stanford University School of Medicine, Stanford, CA, USA.
32. Stanford Cancer Institute, Stanford University School of Medicine, Stanford, CA, USA.
33. Department of Medicine, Division of Hematology, Stanford University School of Medicine, Stanford, CA, USA.
34. Department of Medicine, Division of Cardiovascular Medicine, Stanford University, Stanford, CA, USA.
35. Department of Neurology and Neurological Sciences, Stanford University School of Medicine, Stanford, CA, USA.
36. Division of Gastroenterology, Department of Medicine, University of California, San Francisco, San Francisco, CA, USA.
37. Department of Bioengineering and Therapeutic Sciences, University of California, San Francisco, San Francisco, CA, USA.
38. Department of OB/GYN Stanford University, Palo Alto, CA, USA.
39. Division of Hematology and Oncology, Department of Medicine, Bakar Computational Health Sciences Institute, Institute for Human Genetics, University of California San Francisco, San Francisco, CA, USA.
40. Division of Pediatric Otolaryngology Stanford University School of Medicine, Stanford, CA, USA.
41. Stanford Health Care, Stanford CA, USA.
42. Mass Spectrometry Platform, Chan Zuckerberg Biohub, Stanford, CA, USA.
43. Department of Electrical Engineering and Computer Sciences, University of California Berkeley, Berkeley, CA, USA.
44. Data Intuitive, Flanders, Belgium.
45. Data Mining and Modelling for Biomedicine group, VIB Center for Inflammation Research, Ghent, Belgium.
46. Department of Applied Mathematics, Computer Science, and Statistics, Ghent University, Ghent, Belgium.
47. Department of Computer Science, Columbia University; New York, NY, USA.
48. Department of Biomedicine, Aarhus University, Aarhus, Denmark.
49. Department of Otolaryngology, Vanderbilt University Medical Center, Nashville, TN, USA.
50. Department of Cellular Molecular Pharmacology, University of California, San Francisco, San Francisco, CA, USA.
51. Department of Cell & Tissue Biology, University of California San Francisco, San Francisco, CA, USA.
52. Center for Cancer Research Medical University of Vienna Borschkegasse 8a 1090 Vienna, Austria.
53. Department of Urology, Stanford University School of Medicine, Stanford, CA, USA.
54. Department of Pharmaceutical Chemistry, University of California San Francisco, San Francisco, CA.
55. Division of Hematology/Oncology, Department of Medicine, San Francisco Veterans Affairs Health Care System, San Francisco, CA, USA.
56. Pharmacology and Molecular Biology Departments, Schools of Medicine and Biological Sciences, University of California, San Diego, CA, USA.
57. Department of Pediatrics, Division of Pulmonary Medicine, Stanford University, Stanford, CA, USA.
58. Department of Surgery, Stanford University School of Medicine, Stanford, CA, USA.
59. Veterans Affairs Palo Alto Health Care System, Palo Alto, CA, USA.
60. Department of Computer Science, Stanford University, Stanford, CA.
